# Structure and dynamics of SARS-CoV-2 proofreading exoribonuclease ExoN

**DOI:** 10.1101/2021.04.02.438274

**Authors:** Nicholas H. Moeller, Ke Shi, Özlem Demir, Surajit Banerjee, Lulu Yin, Christopher Belica, Cameron Durfee, Rommie E. Amaro, Hideki Aihara

## Abstract

High-fidelity replication of the large RNA genome of coronaviruses (CoVs) is mediated by a 3′-to-5′ exoribonuclease (ExoN) in non-structural protein 14 (nsp14), which excises nucleotides including antiviral drugs mis-incorporated by the low-fidelity viral RNA-dependent RNA polymerase (RdRp) and has also been implicated in viral RNA recombination and resistance to innate immunity. Here we determined a 1.6-Å resolution crystal structure of SARS-CoV-2 ExoN in complex with its essential co-factor, nsp10. The structure shows a highly basic and concave surface flanking the active site, comprising several Lys residues of nsp14 and the N-terminal amino group of nsp10. Modeling suggests that this basic patch binds to the template strand of double-stranded RNA substrates to position the 3′ end of the nascent strand in the ExoN active site, which is corroborated by mutational and computational analyses. Molecular dynamics simulations further show remarkable flexibility of multi-domain nsp14 and suggest that nsp10 stabilizes ExoN for substrate RNA-binding to support its exoribonuclease activity. Our high-resolution structure of the SARS-CoV-2 ExoN-nsp10 complex serves as a platform for future development of anti-coronaviral drugs or strategies to attenuate the viral virulence.

## Introduction

The 29.9 kb single-stranded RNA genome of SARS-CoV-2, the causative agent of the global COVID-19 pandemic, is replicated and transcribed by the viral RNA-dependent RNA polymerase (RdRp, nsp12) (1–3). Unlike the high-fidelity cellular replicative DNA polymerases, viral RdRp enzymes including the coronavirus (CoV) RdRp do not contain a proofreading exonuclease domain to ensure high fidelity. The resulting higher mutation rate (10^-4^ to 10^-6^ substitutions/nucleotide/round of replication) is generally thought to promote rapid viral adaptation in response to selective pressure (4–6). However, the lack of proofreading activity in RdRp poses a particular challenge for the replication of coronaviruses, which feature the largest known RNA virus genomes (27 ~ 32 kb, up to twice the length as the next-largest non-segmented RNA viral genomes) (7, 8). It has been reported that SARS-CoV nsp12 is the fastest viral RdRp known but with an error rate more than one order of magnitude higher than the generally admitted error rate of viral RdRps (9), clearly necessitating a unique proofreading mechanism.

To mitigate the low fidelity of RdRp, all coronaviruses encode a 3′-to-5′ exoribonuclease (ExoN) in nsp14 (10–12). Mutations of SARS-CoV-2 nsp14 exhibit strong association with increased genome-wide mutation load (13, 14), and genetic inactivation of ExoN in engineered SARS-CoV and murine hepatitis virus (MHV) leads to 15 to 20-fold increases in mutation rates (7, 15, 16). Furthermore, in a mouse model, SARS-CoV with inactivated ExoN shows a mutator phenotype with decreased fitness and lower virulence over serial passage, suggesting a potential strategy for generating a live, impaired-fidelity coronavirus vaccine (17). Alternatively, recent studies show that ExoN inactivation is lethal for SARS-CoV-2 and Middle East Respiratory Syndrome (MERS)-CoV (18), hinting at additional functions for ExoN in viral replication. Indeed, the ExoN activity has been reported to mediate extensive viral RNA recombination required for subgenomic mRNA synthesis during normal replication of CoVs including SARS-CoV-2 (19), and it was shown to be required for resistance to the antiviral innate immune response for MHV (20). ExoN inactivation also significantly increases the sensitivity of CoVs to nucleoside analogs that target RdRp, which is consistent with the biochemical activity of ExoN to excise mutagenic or chain-terminating nucleotides mis-incorporated by RdRp (21–23). These observations combine to suggest that chemical inhibition of ExoN could be an effective antiviral strategy against CoVs. In this study, we determined a high-resolution crystal structure of the SARS-CoV-2 ExoN-nsp10 complex and studied its biochemical activities. Furthermore, we used molecular dynamics (MD) simulations to better understand the dynamics of nsp14, nsp10, and their interaction with RNA.

## Results

The multifunctional SARS-CoV-2 nsp14 consists of the N-terminal ExoN domain involved in proofreading and the C-terminal guanine N7 methyl transferase (N7-MTase) domain that functions in mRNA capping. We co-expressed in bacteria the full-length 527-residue SARS-CoV-2 nsp14 or its N-terminal fragment (residues 1 to 289) containing only the ExoN domain, with full-length 139-residue nsp10 in both cases and purified the heterodimeric complexes. The nsp14-nsp10 and ExoN-nsp10 complexes both showed the expected 3′-to-5′ exonuclease activity on a 5′-fluorescently labeled 20-nucleotide (nt) RNA (LS2U: 5′-GUCAUUCUCCUAAGAAGCU**U**; similar to ‘LS2’ used previously in SARS-CoV ExoN studies (21)) (**Fig. 1A. B**). Although LS2U RNA by itself served as a substrate, more extensive degradation was observed when it was annealed to an unlabeled 40-nt template strand (LS15A_RNA; **Table 1**) to generate a double-stranded (ds) RNA with a 20-nt 5′-overhang. Introducing a base-mismatch at the 3′ end of the degradable strand by using an alternative bottom strand (LS15_RNA; **Table 1**) had no discernable effect on the processing by either complex (**Fig. 1A. B**). When DNA was used as the template strand (LS15_DNA; **Table 1**) to generate an RNA/DNA heteroduplex substrate that is expected to take the A-form conformation similarly to dsRNA, the activity was observed but weaker than for dsRNA. No nuclease activity was observed on a 5′-fluorescently labeled 20-nt DNA (LS2_DNA; **Table 1**), whether the template strand was RNA (LS15_RNA), DNA (LS15_DNA; **Table 1**), or absent. A 20-nt poly-U RNA (U20_RNA; **Table 1**), which is less likely to adopt secondary structures than LS2U, did not serve as a substrate by itself but was degraded extensively when supplemented with a complementary 30-nt poly-A RNA (A30_RNA; **Table 1**) (**Fig. 1C**). Collectively, these results show that the N-terminal ExoN domain of SARS-CoV-2 nsp14 is sufficient for binding to nsp10 to form an active exoribonuclease complex that preferentially degrades dsRNA. For comparison, we also generated a corresponding SARS-CoV ExoN-nsp10 complex, which showed similar activities to SARS-CoV-2 ExoN-nsp10 (**Fig. 1C, Supplementary Fig. 1**).

**Fig. 1.**
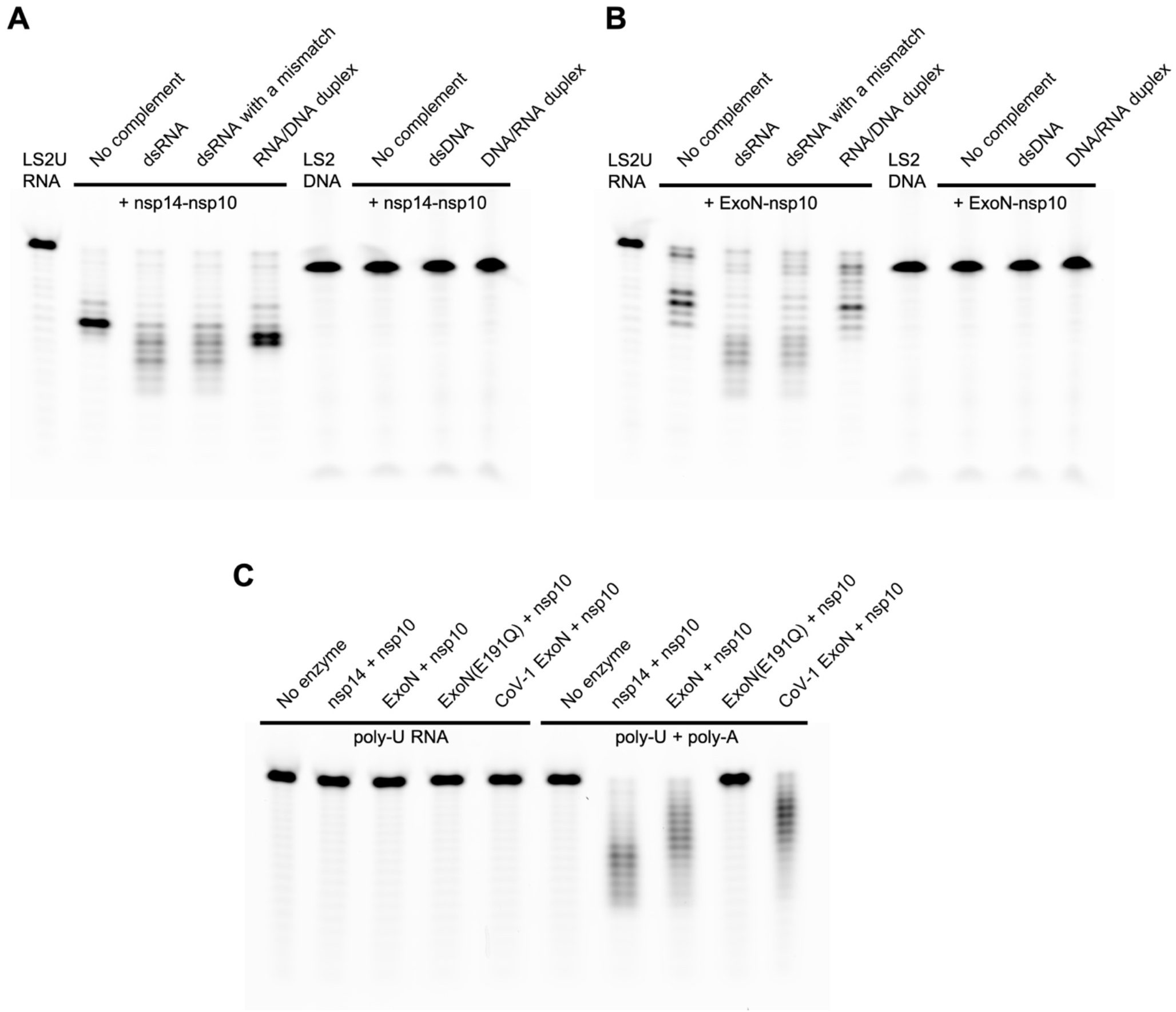
Biochemical activities of nsp14 or its N-terminal ExoN domain, in complex with nsp10. **A**, Exonuclease activities of SARS-CoV-2 full-length nsp14-nsp10 complex on various RNA and DNA substrates. **B**, Exonuclease activities of SARS-CoV-2 ExoN (nsp14 residues 1-289)-nsp10 complex on the same set of RNA and DNA substrates as in (a). **C**, Exonuclease activities of SARS-CoV-2 full-length nsp14-nsp10, SARS-CoV-2 ExoN-nsp10, and SARS-CoV ExoN-nsp10 complexes on poly-U RNA in the absence (left) or presence (right) of unlabeled poly-A RNA. Please see **Table 1** for the substrate sequences.

**Table 1:**
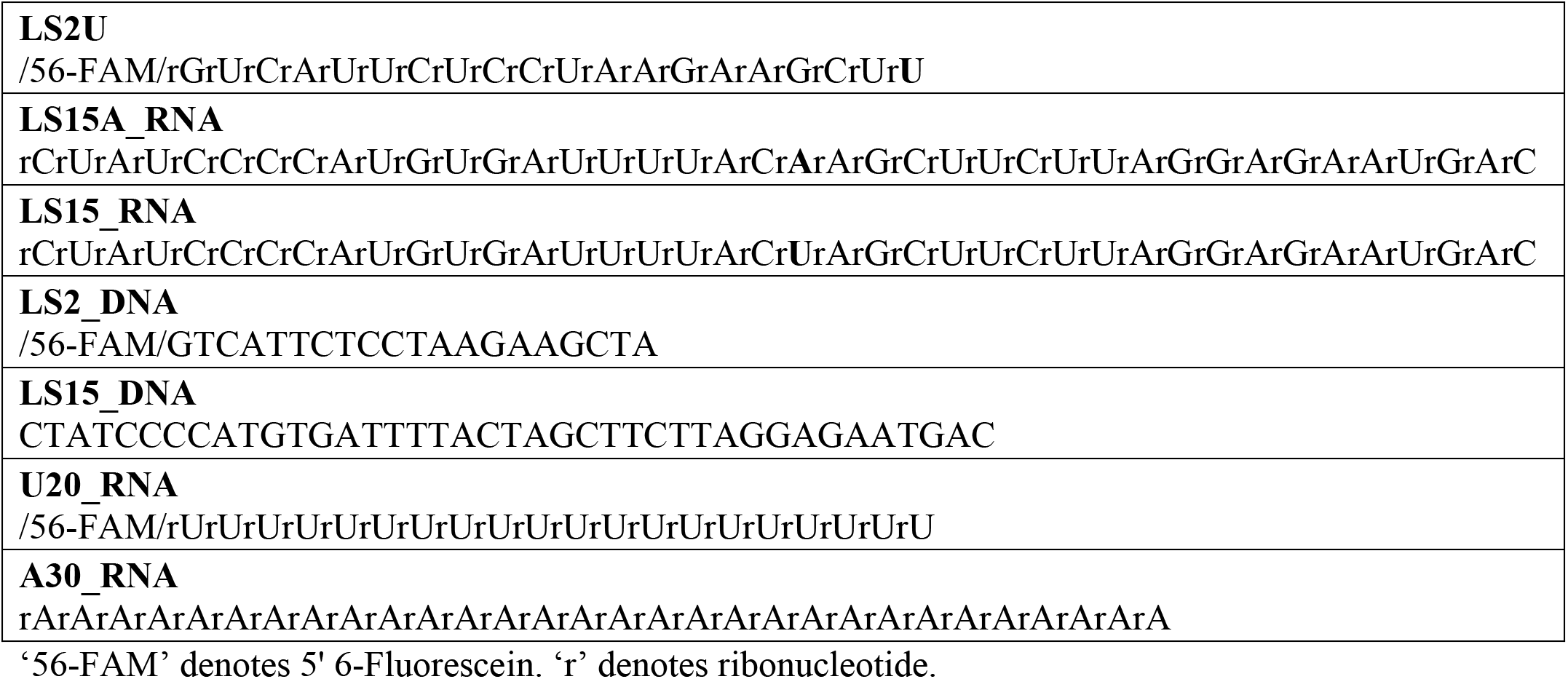
Oligonucleotides used in biochemical assays

Previous X-ray crystallographic studies have provided the structure of SARS-CoV nsp14-nsp10 complex at resolutions ranging from 3.2 to 3.4 Å (21, 24). To obtain higher resolution view of a CoV exoribonuclease complex and to reveal possible structural difference between SARS-CoV and SARS-CoV-2 ExoN, we have crystallized the SARS-CoV-2 ExoN-nsp10 complex. An ExoN variant with a nuclease-inactivating mutation (E191Q) (**Fig. 1C, Supplementary Fig. 1**) was used in our crystallographic studies as it was expressed more robustly and generated a more stable complex with nsp10 than wild-type ExoN. We obtained crystals under two different conditions, one containing ammonium tartrate and the other containing magnesium chloride (MgCl_2_), albeit in the same crystal form. The structures were determined by molecular replacement phasing and refined to 1.64 and 2.10-Å resolution for the tartrate and magnesium-bound crystals, respectively (**Fig. 2A, Table 2**). The final models consist of nsp14 residues Asn3 to Arg289 (Val287 for the lower resolution structure) and nsp10 residues Ala1 to Cys130, with two zinc ions bound to each polypeptide chain. As expected from high sequence conservations, SARS-CoV-2 ExoN-nsp10 complex shows high structural similarity to its counterpart from SARS-CoV (root-mean-square deviation of 0.95 Å for all main chain atoms against 5C8T (24)), whose shape was previously described to resemble ‘hand (ExoN) over a fist (nsp10)’ (21) (**Fig. 2B**). A superposition between the SARS-CoV and SARS-CoV-2 ExoN-nsp10 structures shows only relatively small (3.0 Å or less) deviations in several regions of the complex, including the tip of the ‘fingers’ region of ExoN comprising nsp14 residues 40 ~ 50, and surface-exposed loops of nsp10 (**Supplementary Fig. 2**).

**Fig. 2.**
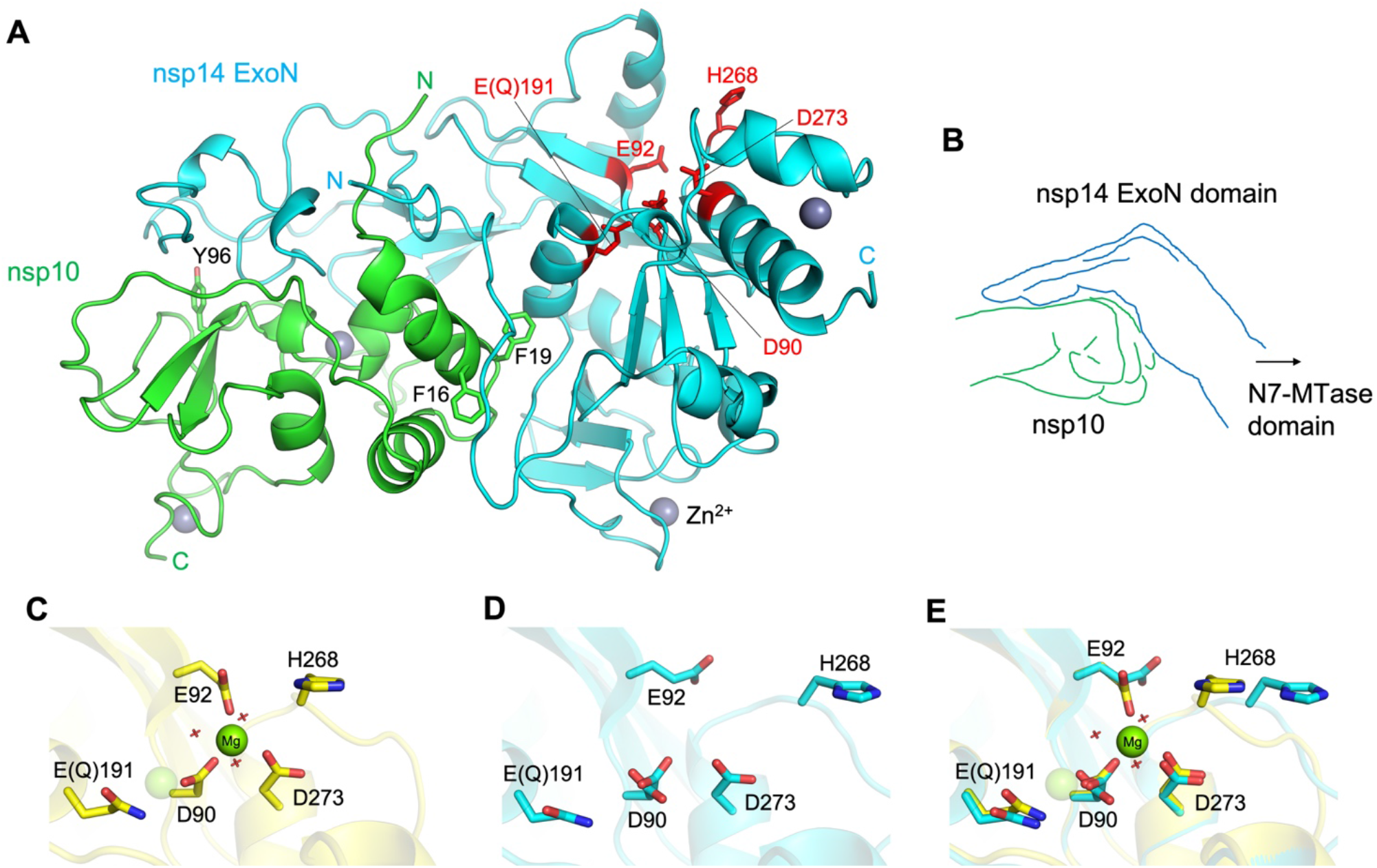
SARS-CoV-2 ExoN-nsp10 structure and its active site flexibility. **A,** Overall structure of nsp14(1-289)-nsp10 complex. The N-terminal ExoN domain of nsp14 is shown in cyan and nsp10 in green. The ExoN active site residues are highlighted as red sticks. Key aromatic residues of nsp10 in the protein-protein interface are also shown as sticks. Gray spheres represent zinc ions. **B**, A schematic illustration of hand (ExoN) over a fist (nsp10). **C**, ExoN active site in the presence of Mg^2+^. The magnesium ion is shown as a solid sphere scaled at half the van der Waals radius. The second Mg^2+^-binding site, indicated by a transparent sphere, is unoccupied in our structure presumably due to the E191Q mutation. **D**, Mg^2+^-free active site as observed in the tartrate-bound crystal. Asp90 side chain shows a dual conformation. **E**, Superposition of **C** and **D**, highlighting the conformational changes upon Mg^2+^-binding.

**Table 2:** Summary of X-ray data collection and model refinement statistics

|  | ExoN-nsp10 (7MC5) | ExoN-nsp10-Mg<sup>2+</sup> (7MC6) |
| --- | --- | --- |
| **Data collection** |  |  |
| Wavelength (Å) | 0.979 | 0.979 |
| Resolution range (Å) | 57.7 - 1.64 (1.70 - 1.64) | 42.6 - 2.10 (2.18 - 2.10) |
| Space group | P2<sub>1</sub>2<sub>1</sub>2<sub>1</sub> | P2<sub>1</sub>2<sub>1</sub>2<sub>1</sub> |
| Unit cell (*a,b,c* in Å) | 63.74 67.48 111.25 | 61.67 70.32 108.54 |
| Total reflections | 258196 (22096) | 105896 (10815) |
| Unique reflections | 58702 (5273) | 27756 (2767) |
| Multiplicity | 4.4 (4.2) | 3.8 (3.9) |
| Completeness (%) | 98.81 (90.43) | 98.25 (99.43) |
| $\langle I/\sigma(I) \rangle$ | 12.57 (1.48) | 10.70 (1.96) |
| R<sub>merge</sub> | 0.148 (1.22) | 0.078 (0.928) |
| R<sub>meas</sub> | 0.166 (1.40) | 0.091 (1.082) |
| R<sub>p.i.m.</sub> | 0.076 (0.660) | 0.045 (0.543) |
| CC<sub>1/2</sub> | 0.995 (0.394) | 0.997 (0.524) |
| **Refinement** |  |  |
| Reflections, working set | 58626 (5273) | 27755 (2768) |
| Reflections, test set | 2826 (251) | 1364 (132) |
| R<sub>work</sub> | 0.166 (0.354) | 0.197 (0.306) |
| R<sub>free</sub> | 0.197 (0.371) | 0.219 (0.346) |
| No. of non-H atoms | 3890 | 3447 |
| Macromolecules | 3264 | 3221 |
| Ligands | 117 | 42 |
| Solvent | 509 | 184 |
| Protein residues | 417 | 415 |
| R.m.s. deviations |  |  |
| Bond length (Å) | 0.011 | 0.001 |
| Bond angles (°) | 1.10 | 0.41 |
| Ramachandran plot |  |  |
| Favored (%) | 96.85 | 96.84 |
| Allowed (%) | 2.91 | 2.92 |
| Outliers (%) | 0.24 | 0.24 |
| Average *B* factor (Å<sup>2</sup>) | 26.61 | 44.43 |
| Macromolecules | 24.60 | 44.11 |
| Ligands | 37.61 | 54.76 |
| Solvent | 36.94 | 47.72 |

Statistics for the highest-resolution shell are shown in parentheses.

While our structures of SARS-CoV-2 ExoN-nsp10 obtained in the two different crystallization conditions are highly similar to each other, they show notable differences in the exonuclease active site located around the ‘knuckles’ of ExoN. In the crystal grown in the presence of MgCl_2_, we observed a magnesium ion octahedrally coordinated by Asp90, Glu92, Asp273, and three water molecules (**Fig. 2C, Supplementary Fig. 3d**). Another magnesium ion required for the conserved two-metal ion mechanism of 3′-5′ editing exonucleases (25, 26) was not observed. The previously reported SARS-CoV nsp14-nsp10 structures also showed only one metal ion, bound at an alternative site between Asp90 and Glu191 (21, 24). This site is unoccupied in our structure presumably due to the E191Q mutation. In contrast, the higher resolution tartrate-bound structure shows a unique configuration of metal-free active site (**Fig. 2D, Supplementary Fig. 3c**). Without the magnesium ion, Asp90 takes two distinct conformers with its carboxylate group in orthogonal orientations. Glu92 is pointed away from Asp90/Asp273 and hydrogen-bonded to Gln108 side chain, whereas His268 in turn is flipped away from Glu92. A comparison between the Mg^2+^-bound and free structures shows a significant rearrangement for residues Gly265 to Val269 including the main chain atoms, accompanying an inward movement of His268 upon Mg^2+^-binding (**Fig. 2E)**. These observations demonstrate high flexibility of the ExoN active site in the absence of divalent metal co-factors.

To obtain an idea about how SARS-CoV-2 ExoN-nsp10 complex engages RNA substrates, we modeled an RNA-bound ExoN-nsp10 structure based on the double-stranded (ds) RNA-bound structures of Lassa virus nucleoprotein (NP) exonuclease domain, which is another DEDDh-family 3′-to-5′ exoribonuclease. A superposition of the Lassa NP-RNA complex (27, 28) on ExoN-nsp10 based on their conserved catalytic residues (Lassa NP: D389/E391/D466/H528/D533 according to the numbering in 4FVU (27), vs. SARS-CoV-2 ExoN: D90/E92/E191/H268/D273) places the A-form dsRNA in a shallow groove on ExoN surface adjacent to the active site, with remarkable shape complementarity (**Fig. 3 B, C**). In this model, the sugar-phosphate backbone of the non-degradable (template) RNA strand tracks a positively charged patch on the ExoN surface including Lys9 and Lys61, whereas the 3′ end of its complementary (degradable) strand is presented to the active site. The extensive protein contacts made by the non-degradable strand in a dsRNA substrate is consistent with the preference for dsRNA substrates by SARS-CoV-2 ExoN as shown above (**Fig. 1**) and by SARS-CoV ExoN reported earlier (29). Notably, we observed ordered tartrate ions from the crystallization condition bound to this basic patch in our crystal structure, potentially mimicking RNA backbone phosphate interactions (**Supplementary Figs. 3a, 3b,** and **4**).

**Fig. 3.**
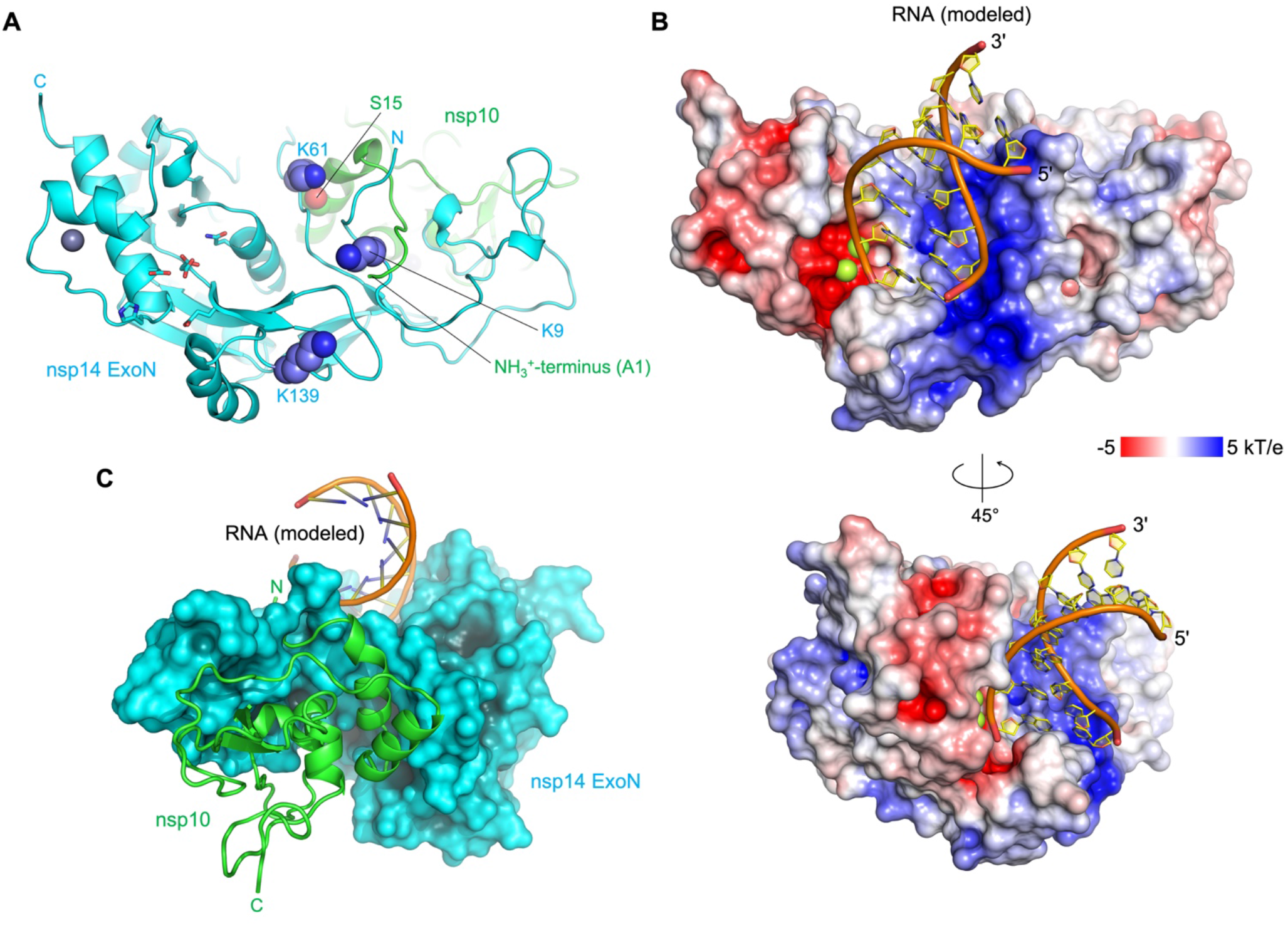
Location of the basic patch and an ExoN-nsp10-RNA complex model. **A,** Locations of the ExoN lysine residues forming the basic patch. Note that Lys9 and Lys61 interact with the N-terminus (Ala1) and Ser15 of nsp10, respectively. **B**, A hypothetical model of ExoN-nsp10-dsRNA complex, viewed from two different orientations. The protein surface is colored according to the electrostatic potential calculated using APBS (46). **C**, Backside of the ExoN-nsp10-dsRNA model, viewed from the ExoN-nsp10 interface. Nsp10 is shown as green ribbon.

Our hypothetical model described above suggests that the basic patch of ExoN helps position the substrate RNA for exonucleolytic degradation. Lys9 and Lys61 are involved in the RNA backbone interaction in our model. In addition, Lys139 is located farther down along the basic patch toward the direction of the 5′-overhang of the template strand (**Fig. 3A**). Thus, we tested the activities of SARS-CoV-2 ExoN with single amino acid substitutions, K9A, K61A, and K139A. These ExoN mutants were co-expressed with nsp10 and purified as heterodimeric complexes. In the exoribonuclease assay using the RNA substrates described above, all three lysine-to-alanine mutants showed lower activity than wild-type ExoN (**Fig. 4**). In particular, the K9A and K61A substitutions caused severer defect than K139A, consistent with our dsRNA-binding model (**Fig. 3 B, C**). While the precise conformation of LS2U RNA in the absence of a complementary strand is unknown, its binding to ExoN must also depend on these Lys residues, underscoring the importance of electrostatic interactions with RNA by the mutated lysine residues in the ExoN activity.

**Fig. 4.**
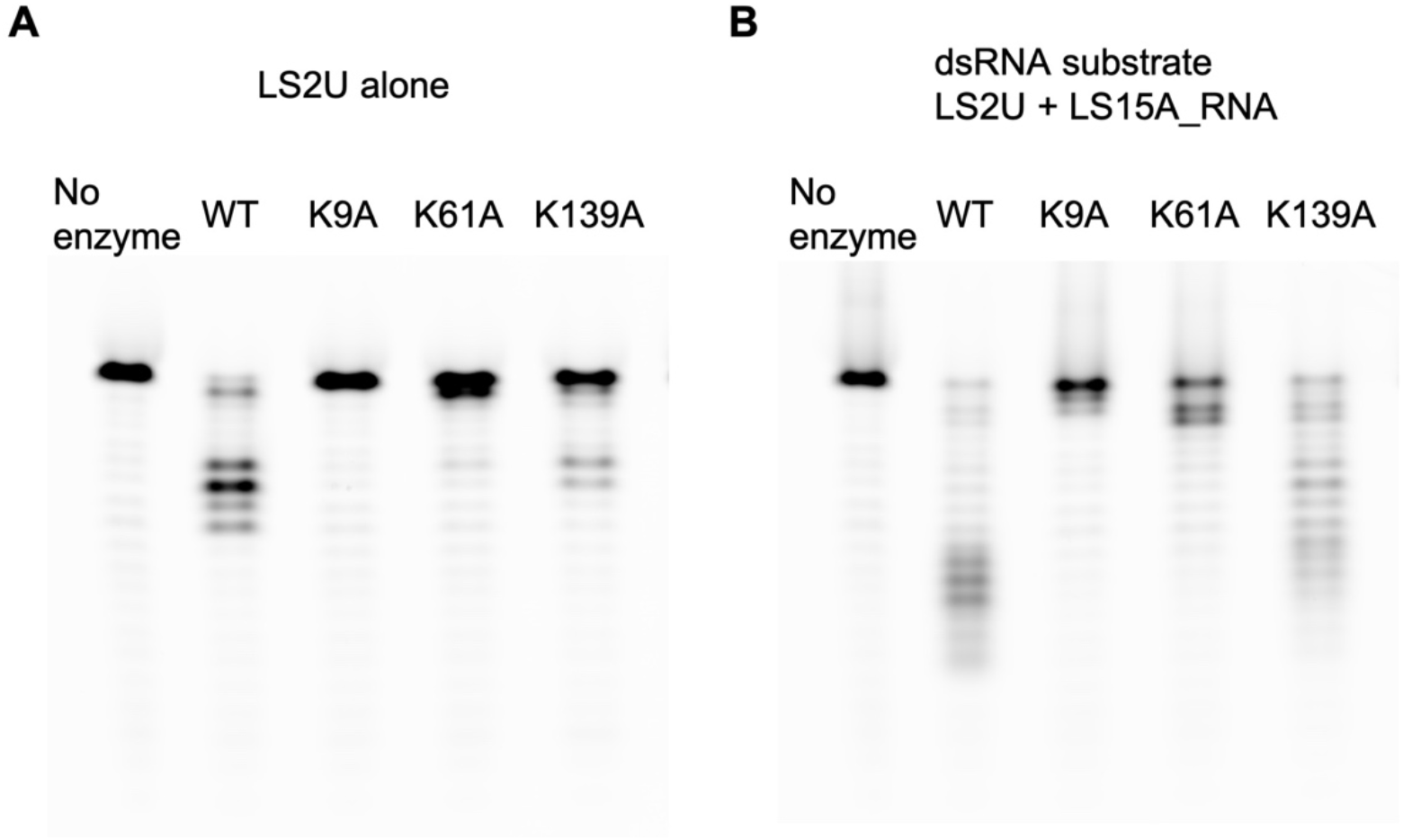
Activities of ExoN lysine mutants. Exoribonuclease activities of SARS-CoV-2 ExoN-nsp10 complex and its lysine-to-alanine point mutant derivatives. **A**, Processing of LS2U RNA without a complementary strand. **B**, LS2U RNA annealed with the fully complementary LS15A RNA (dsRNA substrate). Please see **Table 1** for the substrate sequences.

Previous studies showed that the exoribonuclease activity of nsp14 is strongly stimulated by nsp10 for both SARS-CoV and SARS-CoV-2 (29–32). In our crystal structure, the N-terminal residues of ExoN and those of nsp10 are wrapped around each other in a ‘criss-cross’ arrangement and forming several hydrogen-bond contacts, including one between nsp14 Lys9 and nsp10 Ala1 (**Supplementary Fig. 3a**). In addition, the first a-helix of nsp10 interacts with the ExoN loop harboring nsp14 Lys61, where the main chain amide group of Lys61 is hydrogen-bonded to the side chain of nsp10 Ser15 (**Fig. 3A**). In the absence of nsp10 supporting the RNA-binding groove from the back (**Fig. 3C, Supplementary Figs. 5, 6**), the N-terminal residues of ExoN including nsp14 Lys9 and those around Lys61 are likely to be more flexible. Moreover, the terminal amino group of nsp10 Ala1 is part of the basic patch and involved in direct RNA backbone contact in our protein-RNA docking model (**Fig. 3B, Supplementary Fig. 5**). These observations may together explain the strong stimulation of ExoN activity by nsp10.

To obtain further insights into the role of nsp10 and to support our RNA-binding model, we performed explicitly solvated, all-atom molecular dynamics (MD) simulations of full-length SARS-CoV-2 nsp14, constructed from our ExoN-nsp10 co-crystal structure and a homology model of the C-terminal N7-MTase domain. Three independent copies of MD simulations totaling 2.6-μs were performed for each of nsp14 alone, nsp14-nsp10 complex, and the nsp14-nsp10-RNA complex based on our docking model described above. In addition, three independent copies of Gaussian-accelerated MD simulations (GAMD) totaling 0.6-μs were performed for each system to enhance conformational sampling. Comparing trajectories of these simulations for the 3 systems, the most noticeable difference is an extreme flexibility of the ‘fingers’ region of ExoN primarily comprising its N-terminal residues (nsp14 residues 1-60), which showed large deviations from the starting model and eventually became highly disordered in the absence of nsp10. A principal component analysis for the 3 systems show that the conformational space sampled by nsp14 is significantly larger in the absence of nsp10 (**Fig. 5A, Supplementary Fig. 7**).

**Fig. 5.**
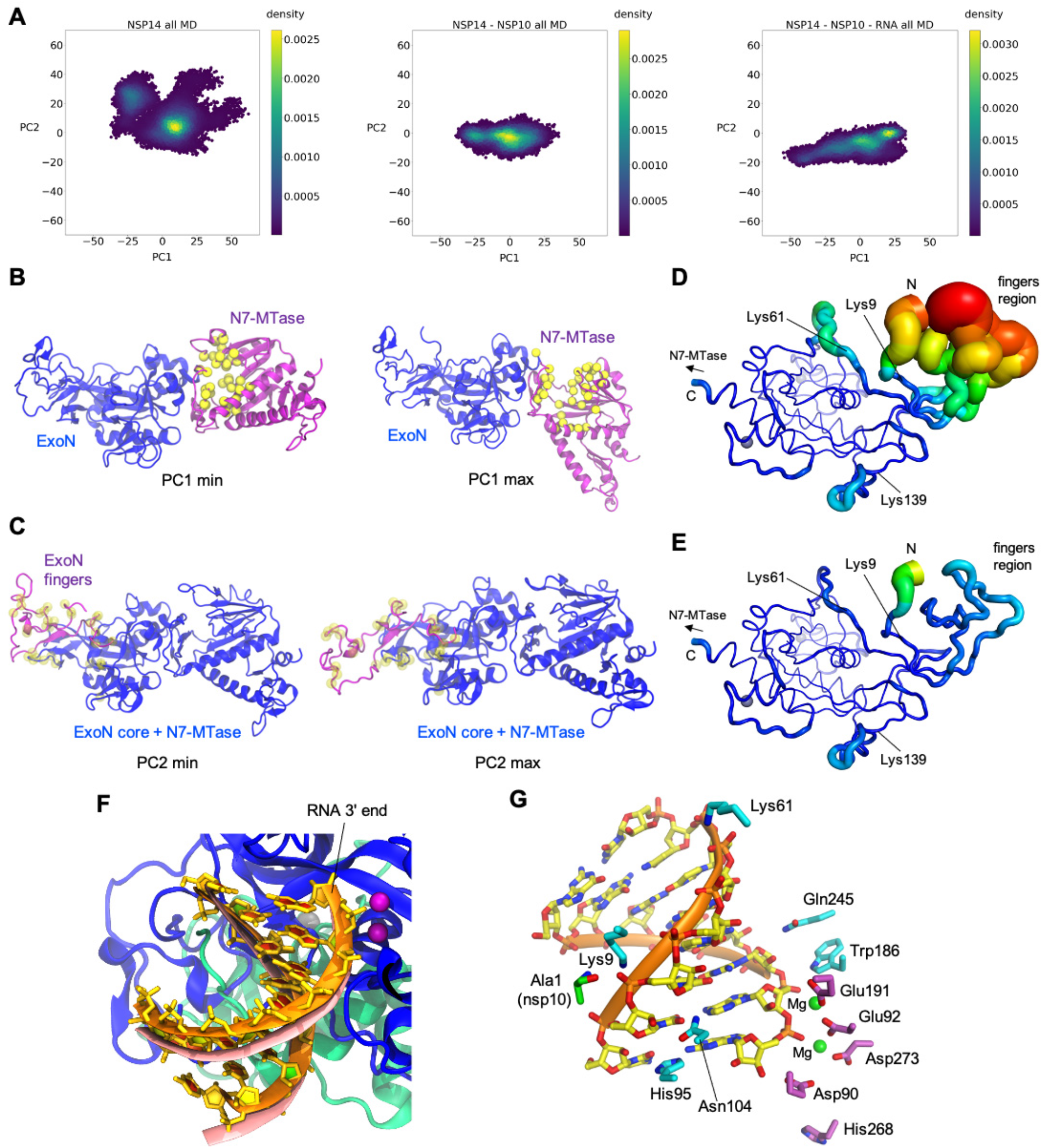
MD simulations. **A,** Principal component analysis depicting differential conformational sampling for the 3 systems in MD simulations. **B,** Structures that correspond to PC1 minimum and maximum values for the nsp14-alone system. N7-MTase and ExoN domains of nsp14 are depicted in purple and blue ribbons, respectively. Yellow spheres represent the Ca atoms of residues that constitute the binding site of SAM and GpppA substrates of N7-MTase based on homology to SARS-CoV nsp14 N7-MTase crystal structures (PDB ID: 5C8S and 5C8T)(24). **C,** Structures that correspond to PC2 minimum and maximum values for the nsp14-alone system. N-terminal region (residues 1-71) of nsp14 is depicted in purple ribbons while the rest of nsp14 is depicted in blue ribbons. Transparent yellow spheres represent the Ca atoms of nsp14 residues that constitute nsp10 binding site. **D,** ExoN domain in nsp14-alone system with root-mean-square fluctuations (RMSF) of Ca atoms depicted on the structure with varying tube thickness and color (low in blue to high in red). The view is similar to that in **Fig. 3A**. **E,** ExoN domain of nsp14-nsp10 system with Ca RMSF depicted on the structure with varying tube thickness and color. **F,** RNA after 1 μs MD simulation (in orange ribbons) of nsp14-nsp10-RNA system superimposed onto RNA of the starting model (salmon). Nsp14 and nsp10 are depicted as blue and green ribbons, respectively. Dark purple spheres represent two Mg ions in the active site. **G,** RNA after 1 μs MD simulation of the nsp14-nsp10-RNA system, with nsp14 ExoN domain (cyan) or nsp10 (green) residues making persistent hydrogen-bond or salt bridge interactions with RNA in MD simulations shown as sticks. The active site residues of ExoN are also shown (purple sticks) with two Mg^2+^ ions as green spheres.

The first principal component (PC1), which is broadly sampled by all 3 systems, corresponds to a large hinge motion of the N7-MTase domain (~50 Å translocation at the distal end, **Supplementary Fig. 8, Supplementary animation 1**). In the conformation with minimal PC1 (**Fig. 5B, left**), the substrates (S-adenosyl methionine [SAM] and GpppA)-binding cleft of the N7-MTase domain abuts against the ExoN domain, leading to occlusion of the substrates. On the other extreme with maximal PC1, the cleft is more open to the solvent (**Fig. 5B, right**). The second principal component (PC2) corresponds to an ordered-to-disordered transition of the ‘fingers’ region of ExoN, which shows a large population of disordered conformations only for the nsp14-alone system as mentioned above (**Fig. 5C, Supplementary Fig. 9, Supplementary animations 2, 3**). Although folding of the core of the ExoN domain does not depend on nsp10, residues Lys9 and Lys61 important for RNA-binding and the surrounding residues show increased flexibility in the absence of nsp10, confirming our prediction above (**Table 3, Fig. 5 D, E, Supplementary Fig. 9, Supplementary animation 3**). The dsRNA molecule in the nsp14-nsp10-RNA complex was stable throughout the simulation with direct RNA phosphate contacts by nsp14 Lys9, Lys61, and the terminal amino group of nsp10 maintained, providing further support for our model for dsRNA-binding (**Fig. 5 F, G, Supplementary Fig. 10**). An ionic interaction between Ala1 of nsp10 and RNA backbone phosphate was particularly persistent and observed for 97 % of the time during the simulations (3.2 Å distance cutoff), which led to a significant stabilization of this residue in the presence of RNA (**Table 3**). Lastly, it is also worth noting that an analysis of the internal dynamics of the N7-MTase domain indicates several highly mobile regions, including loops (residues 289-300, 355-362) flanking the substrates-binding cleft and a loop (residues 454-470) adjacent to the third zinc finger motif of nsp14 distal to the N7-MTase active site (**Supplementary Fig. 11**).

**Table 3:** Root-mean-square fluctuations (RMSF, in Å) of the catalytic residues and RNA-binding residues in the 3 simulated systems. RMSF of Ca atoms were calculated after aligning trajectories to the initial model with respect to Ca atoms of residues 71-289 (core of the ExoN domain). RMSF of all atoms for each residue is presented in parenthesis. Catalytic residues of ExoN are underlined.

|  | **nsp14** | **nsp14-nsp10** | **nsp14-nsp10-RNA** |
| --- | --- | --- | --- |
| **<u>D90</u>** (nsp14) | 0.44 (0.56) | 0.40 (0.52) | 0.35 (0.39) |
| **<u>E92</u>** (nsp14) | 0.61 (1.18) | 0.63 (1.16) | 0.39 (0.76) |
| **<u>E191</u>** (nsp14) | 0.58 (0.76) | 0.56 (0.75) | 0.38 (0.61) |
| **<u>H268</u>** (nsp14) | 1.75 (2.33) | 1.66 (2.21) | 1.34 (2.05) |
| **<u>D273</u>** (nsp14) | 0.58 (0.97) | 0.60 (0.92) | 0.39 (0.45) |
| **K9** (nsp14) | 1.80 (2.59) | 1.16 (1.62) | 0.55 (0.67) |
| **K61** (nsp14) | 2.81 (3.49) | 1.60 (2.26) | 0.73 (1.25) |
| **K139** (nsp14) | 0.95 (1.56) | 0.86 (1.52) | 0.62 (1.18) |
| **A1** (nsp10) |  | 4.04 (4.12) | 0.75 (0.82) |

## Discussion

Our X-ray crystallographic, biochemical, and computational analyses shed light on the substrate preference, structure, and dynamics of the SARS-CoV-2 ExoN-nsp10 exoribonuclease complex and further identified important roles of nsp10 in RNA substrate binding. It is particularly notable that the ExoN-nsp10 complex preferentially degrades dsRNA substrates. This is in contrast to the proofreading exonuclease domain of high-fidelity DNA polymerases, whose active site engages the single-stranded DNA 3′ end in partially melted double-stranded substrates (25, 33), and suggests a unique mechanism of proofreading. The extensive ExoN/nsp10 interface buries a total of 2203 Å^2^ of surfaces from both proteins, spanning both the ‘fingers’ and ‘palm’ regions of ExoN. Folding of the fingers region depends on its interaction with nsp10, which involves several critical residues including nsp10 Tyr96 (31) (**Fig. 2A, Supplementary Fig. 6**). On the other hand, an interesting feature for the interaction in the palm region includes the insertion of Phe16 and Phe19 from the first a-helix of nsp10 into a deep hydrophobic pocket of ExoN, which is essential for the stable complex formation (31). Notably, this hydrophobic pocket is located on the backside from the ExoN active site, where nsp10 Phe19 side chain makes van der Waals contacts with the main chain of an ExoN a-helix harboring one of the catalytic residues Glu191 (**Supplementary Fig. 6**). Thus, targeting said pocket of ExoN by small molecules to block its interaction with nsp10 or potentially to allosterically modulate its catalytic activity could be a possible strategy of inhibition.

MD simulations revealed remarkable flexibility in full-length nsp14 (**Supplementary Figs. 7, 8,** and **Supplementary animations 1, 2**), which affects solvent accessibility of the SAM/GpppA-binding cleft and may play an important role in the catalytic cycle of N7-MTase (**Fig. 5B**). Similar conformational variation, albeit with a much smaller magnitude, was previously observed between two SARS-CoV nsp14 molecules in the asymmetric unit of a crystal (**Supplementary Fig. 8**) (21). Although this hinge motion was observed for all 3 systems (nsp14-alone, nsp14-nsp10, and nsp14-nsp10-RNA) in our simulations, they showed different distributions of the PC1 value (**Supplementary Fig. 7**). In addition, conformational sampling in the nsp14-alone system shows several clusters with distinct combinations of PC1 and PC2 values (**Fig. 5A, left**), suggesting that there may be a long-range interaction between the N-terminal fingers region of ExoN and the C-terminal N7-MTase domain. These observations are consistent with earlier studies showing that single amino acid substitutions R84A and W86A within the ExoN domain completely abolished, while a deletion of the N-terminal 61 residues significantly enhanced, the N7-MTase activity of SARS-CoV nsp14 (34). These mutations in ExoN may have modulated the PC1 motion of nsp14 to affect its N7-MTase activity. Conversely, although we showed in this study that the N7-MTase domain is not essential for the exoribonuclease activity of nsp14 *in vitro*, SARS-CoV nsp14 N7-MTase residues Tyr498 and His487 were shown to be required for RdRp/nsp12 binding (21), which is presumably important in proofreading. Thus, it is likely that the ExoN and N7-MTase domains are functionally dependent on each other, where proper dynamics may be key to support their respective activities and possible coordination. We hope that our structural and functional studies will help future development of ExoN inhibitors to impede the replication of SARS-CoV-2 and related coronaviruses.

### Methods

#### Protein expression and purification

SARS-CoV-2 (GenBank: MN908947.3) nsp14 and nsp14(1-289) were co-expressed with nsp10 in *E. coli* strain BL21(DE3) under the control of T7 promoters. To facilitate purification, a 6xHis tag was added to the N-terminus of nsp14 and nsp14(1-289) with a human rhinovirus (HRV) 3C protease cleavage site. A methionine residue was added to nsp10 to enable translation. Transformed bacteria were cultured in LB medium at 37 °C to the mid-log phase, induced with 0.5 mM and 50 μM (final concentrations) of Isopropyl β-D-1-thiogalactopyranoside and zinc chloride, respectively, and further incubated at 18 °C overnight before being pelleted by centrifugation. Collected bacteria were disrupted by the addition of hen egg white lysozyme and sonication in 20 mM Tris-HCl, pH 7.4, 0.5 M NaCl, 5 mM β-mercaptoethanol, and 5 mM imidazole. The lysate was cleared by centrifugation at 63,000 x g for 1 hour at 4 °C, after which the protein complex in the supernatant was captured by nickel-affinity chromatography and eluted by a linear gradient of imidazole. Eluted proteins were digested with HRV 3C protease overnight at 4 °C, concentrated by ultrafiltration, and passed through a Superdex75 size-exclusion column operating with the same buffer as above except not containing imidazole. The nsp14-nsp10 complexes eluted as a heterodimer were concentrated by ultrafiltration and frozen in small volume aliquots in liquid nitrogen for storage at −80 °C. The ExoN mutant derivatives were generated by site-directed mutagenesis and purified using the same procedure. The protein concentrations were determined based on UV absorbance at 280 nm measured on a Nanodrop8000 spectrophotometer and theoretical extinction coefficients calculated from the protein amino acid sequences.

#### Crystallization and structure determination

Purified nsp14(1-289, E191Q)-nsp10 complex (**Supplementary Fig. 12**) at 17 mg ml^-1^ was crystallized using the hanging drop vapor diffusion method, by mixing the protein solution with an equal volume of reservoir solution including either 0.2 M di-ammonium tartrate, pH 7.0, 20 % polyethylene glycol (PEG) 3,350 (condition 1), or 0.1 M MgCl_2_, 0.1 M Tris-HCl pH 8.5, 20 % PEG 4,000 (condition 2). Both conditions produced thin needles crystals. The crystals were cryo-protected with ethylene glycol and flash-cooled by plunging in liquid nitrogen. X-ray diffraction data were collected at the Northeastern Collaborative Access Team (NE-CAT) beamlines of the Advanced Photon Source (Lemont, IL) and processed using XDS (35). The structure of the SARS-CoV-2 ExoN-nsp10 complex was determined by molecular replacement phasing by PHASER (36), using the crystal structures of SARS-CoV nsp14-nsp10 complex (PDB ID: 5C8T) (24) as the search model. Iterative model building and refinement were performed using COOT (37) and PHENIX (38), respectively. A summary of data collection and model refinement statistics is shown in **Table 2**. Structure images were generated using PyMOL (https://pymol.org/).

#### Exonuclease activity assays

The 5′-fluorescein labeled oligonucleotides (**Table 1**) at 750 nM, in the presence or absence of equimolar complementary unlabeled strands, were incubated with 50 nM nsp14 (or its ExoN domain alone)-nsp10 complexes in 42 mM Tris-HCl, pH 8.0, 0.94 mM MgCl_2_, 0.94 mM dithiothreitol, and 0.009 % Tween-20. After incubation at 37 °C for 10 min, the reactions were stopped by the addition of formamide to 67 % and heating to 95°C for 10 min. The reaction products were separated on a 15 % TBE-Urea gel, which was scanned on a Typhoon FLA 9500 imager.

#### Molecular dynamics simulations

A homology model of full-length SARS-CoV-2 nsp14 was generated for sequence of YP_009725309.1 and taking SARS-CoV nsp14 crystal structure (PDB ID: 5NFY) (21) as a template in Schrödinger Prime module (39). The nsp14 ExoN domain of the homology model was then replaced with the crystal structure of SARS-CoV-2 nsp14 ExoN in complex with nsp10 obtained in this study. E191Q mutation in the crystal structure was reverted computationally to the wild type. For a nsp14-nsp10-RNA model, RNA was modeled based on Lassa NP-RNA complex (PDB ID: 4FVU) (27) and the second Mg ion at the active site was modeled based on a Mn^2+^ ion found in Lassa NP-RNA complex (PDB ID: 4GV9) (28). Three systems were prepared from this model: 1. Full-length nsp14 alone, 2. Full-length nsp14-nsp10 complex, 3. Full-length nsp14-nsp10-RNA complex. Protonation states of titratable amino acids were determined using PropKa analysis (40). Each of these systems were explicitly solvated in TIP3P water box and ions were added to achieve 0.2 M salt concentration. Amber ff14SB (41) and RNA.OL3 force fields are used for protein and RNA, respectively. For zinc ions and zinc-coordinating residues, Cationic Dummy Atom (CADA) parameters were used (42). Conventional MD simulations (cMD) were performed with NAMD2.14 program (43), while Gaussian-accelerated MD simulations (GAMD) were performed with Amber20 program (44). First, each system was minimized in 4 consecutive steps by gradually decreasing restraints. Subsequently, each system was heated from 0 to 310 K slowly, and then equilibrated for about 1 ns by gradually decreasing restraints in 3 consecutive steps. For cMD, three independent copies (2x 1 μs and 1x 0.6 μs) of simulation were run for each system. For GAMD, three independent copies of 0.2 μs of simulation were run for each system using dual boost method following a 20-ns MD run to calculate parameters for GAMD production runs. All cMD and GAMD simulations were performed at 310 K and 1 atm and with a 2 fs timestep. For each system, 32,000 data points with 0.1 ns intervals were collected from simulations and analyzed. Stability of MD simulations are shown with RMSD plots of nsp14 ExoN domain (**Supplementary Fig. 13**). MDTraj (45) was used for some of the MD trajectory analysis.

## Supporting information

Supplementary animations

## Acknowledgements

We thank Daniel Harki and Reuben Harris for thoughtful comments. This work was supported by grants from the US National Institutes of Health (NIGMS R35-GM118047 to H.A., R01-GM132826 to R.E.A., and NCI P01-CA234228 to R.E.A. and H.A.), NSF RAPID MCB-2032054, an award from the RCSA Research Corp., and a UC San Diego Moores Cancer Center 2020 SARS-COV-2 seed grant to R.E.A. This work is based upon research conducted at the Northeastern Collaborative Access Team beamlines, which are funded by the US National Institutes of Health (NIGMS P30 GM124165). The Pilatus 6M detector on 24-ID-C beamline is funded by a NIH-ORIP HEI grant (S10 RR029205). This research used resources of the Advanced Photon Source, a U.S. Department of Energy (DOE) Office of Science User Facility operated for the DOE Office of Science by Argonne National Laboratory under Contract No. DE-AC02-06CH11357 and those of the Minnesota Supercomputing Institute. We are grateful for the efforts of the Texas Advanced Computing Center (TACC) Frontera team and for the compute time made available through a Director’s Discretionary Allocation (made possible by the National Science Foundation award OAC-1818253).

## Data Availability

The atomic coordinates and structure factors for the SARS-CoV-2 ExoN-nsp10 complex structures have been deposited in the RCSB Protein Data Bank, with the accession codes 7MC5 and 7MC6.

---

**Supplementary Fig. 1.**
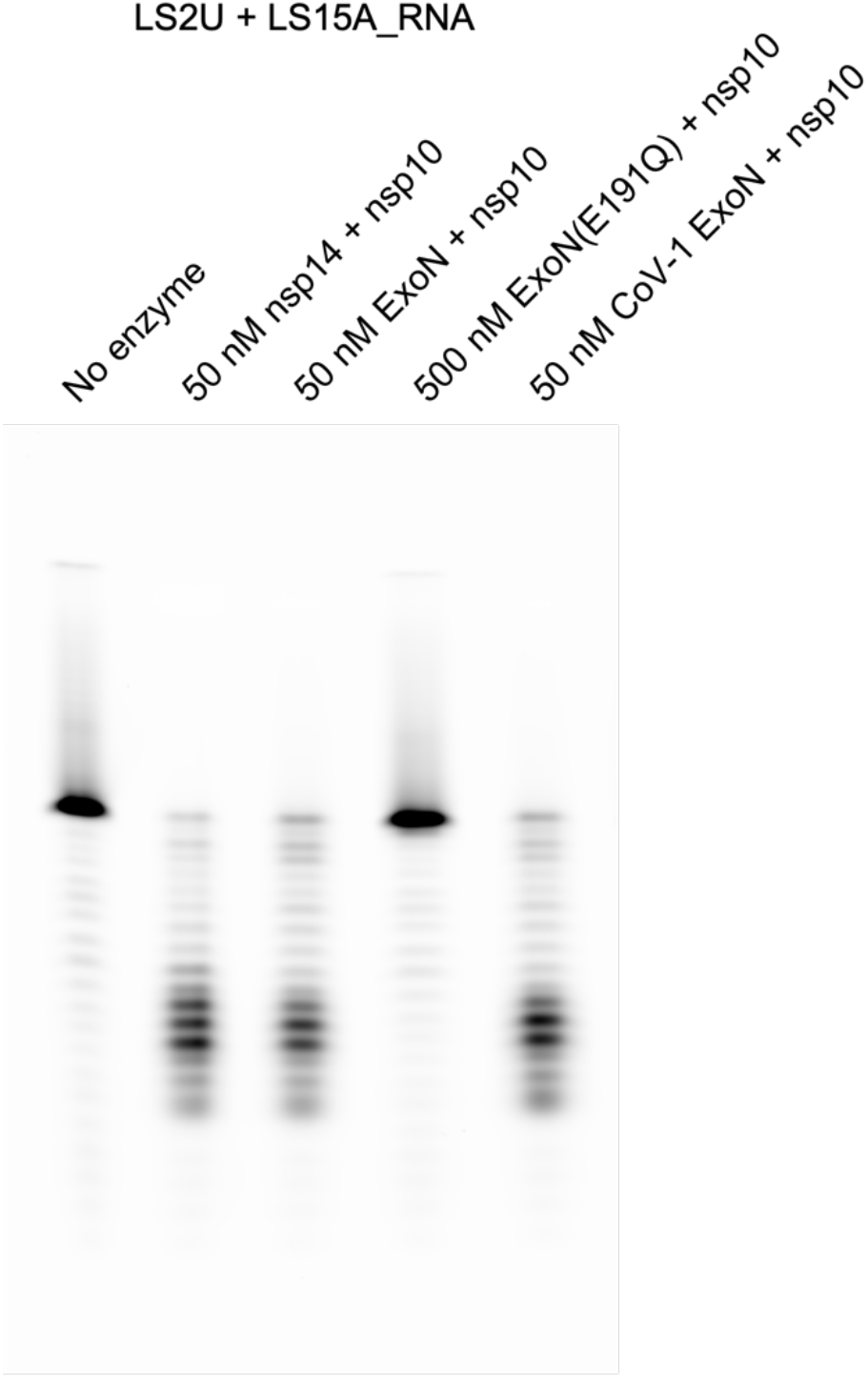
Comparison of exoribonuclease activities. Exoribonuclease activities of SARS-CoV-2 nsp14-nsp10, ExoN-nsp10, and SARS-CoV ExoN-nsp10 complexes on a double-stranded RNA substrate. The inactive E191Q mutant enzyme was tested at a 10 times higher protein concentration.

**Supplementary Fig. 2.**
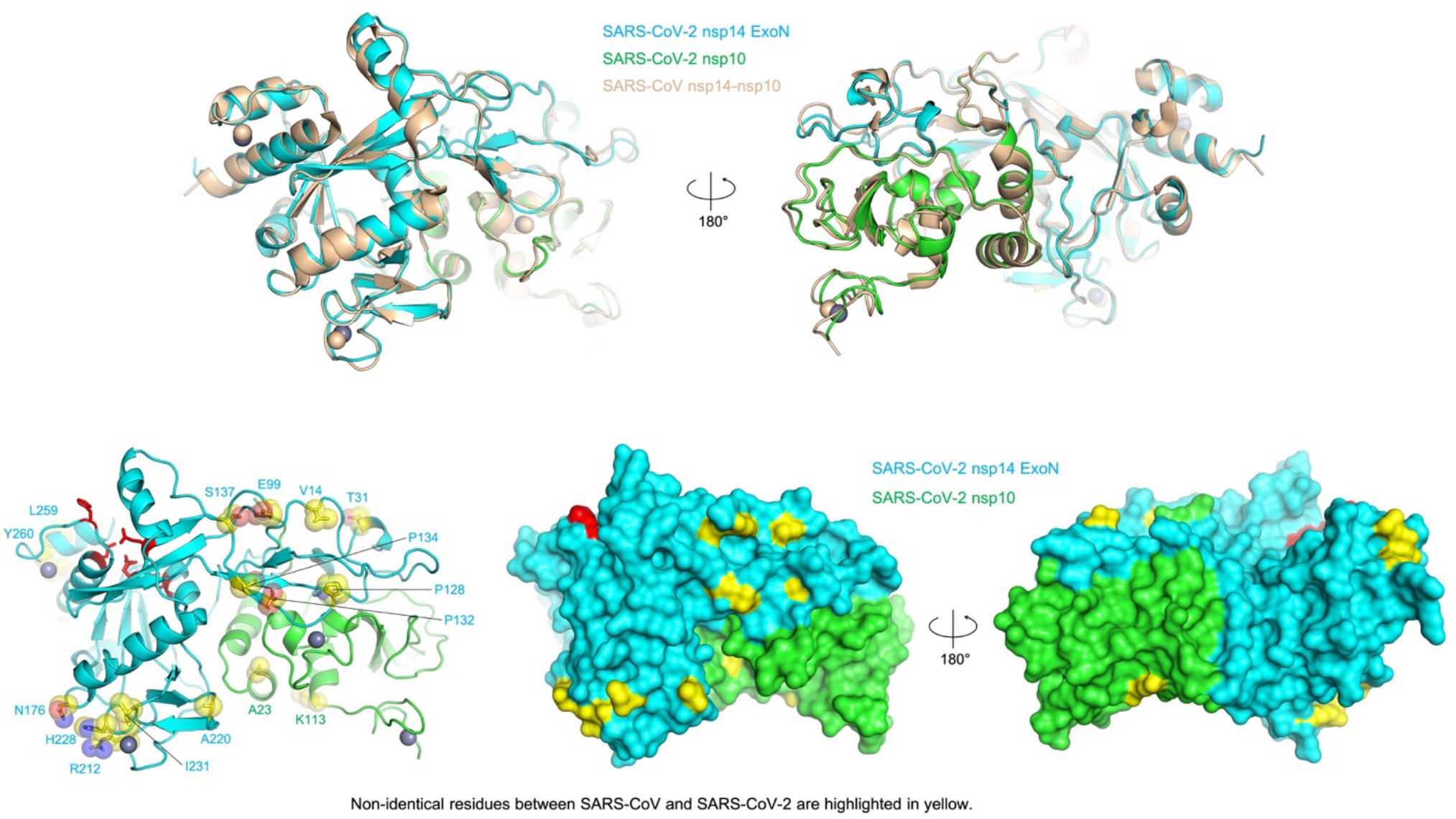
SARS-CoV vs. SARS-CoV-2 ExoN-nsp10 structure comparison. **Top**, A superposition between SARS-CoV (PDB ID: 5C8T) (24) and SARS-CoV-2 (this study) ExoN-nsp10 structures. **Bottom**, Difference in the amino acid sequence between SARS-CoV and SARS-CoV-2 mapped on the ExoN-nsp10 structure and highlighted in yellow. The active site residues are shown in red.

**Supplementary Fig. 3.**
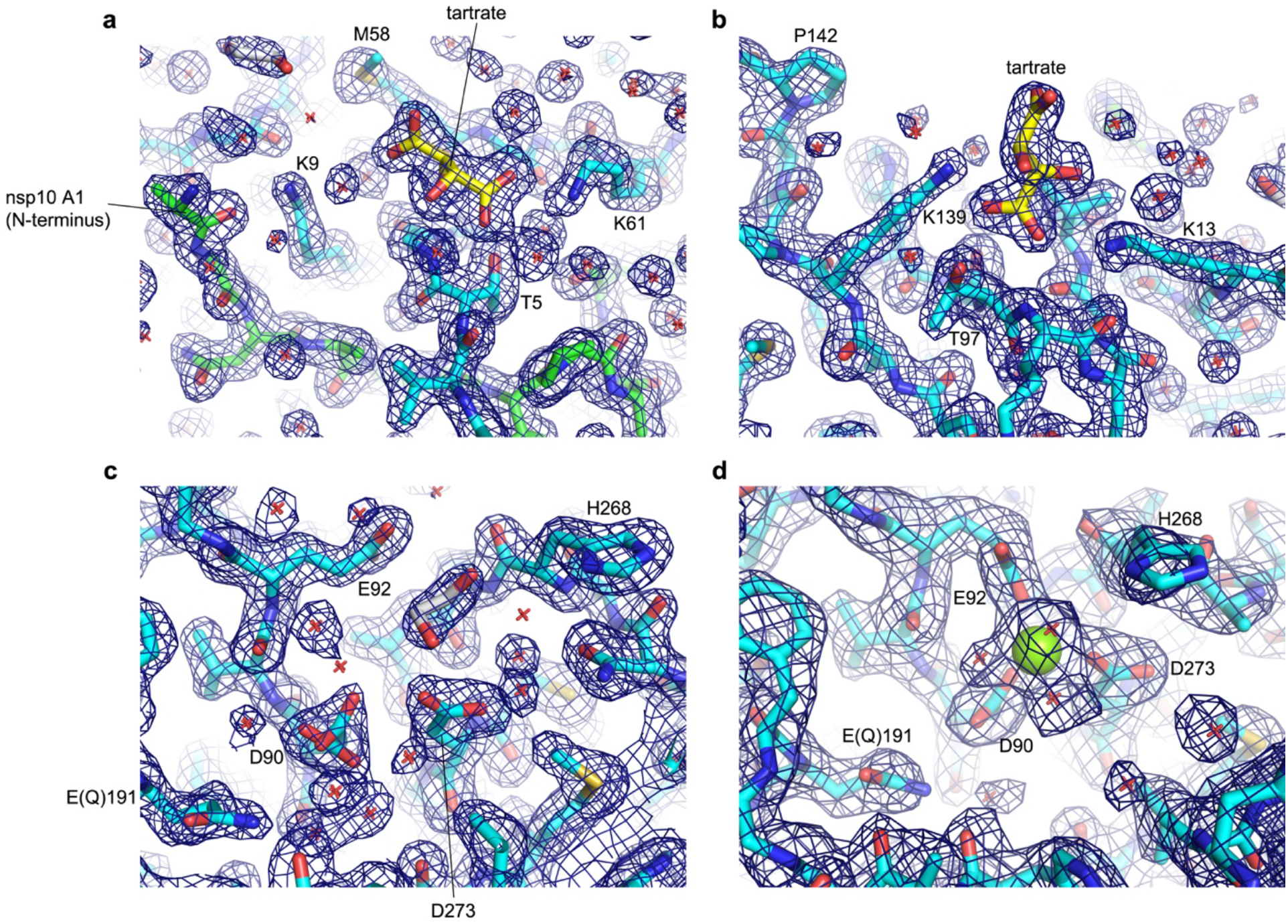
Electron density maps. 2mFo-DFc map contoured at 1.0 σ is shown for the higher resolution (1.64 Å) tartrate-bound structure in **a-c**, and for the lower resolution (2.10 Å) Mg^2+^-bound structure in **d**. **a**, Region including Lys9 and Lys61 of nsp14/ExoN and the N-terminus of nsp10 (The crystallized protein has additional methionine residue on the N-terminus, which is likely to be disordered) with a bound tartrate molecule. **b**, Region including Lys139 and Lys13 with a tartrate molecule bound between the two lysine side chains. **c**, Mg^2+^-free active site. An ethylene glycol molecule used as the cryo-protectant was observed. **d**, Mg^2+^-bound active site.

**Supplementary Fig. 4.**
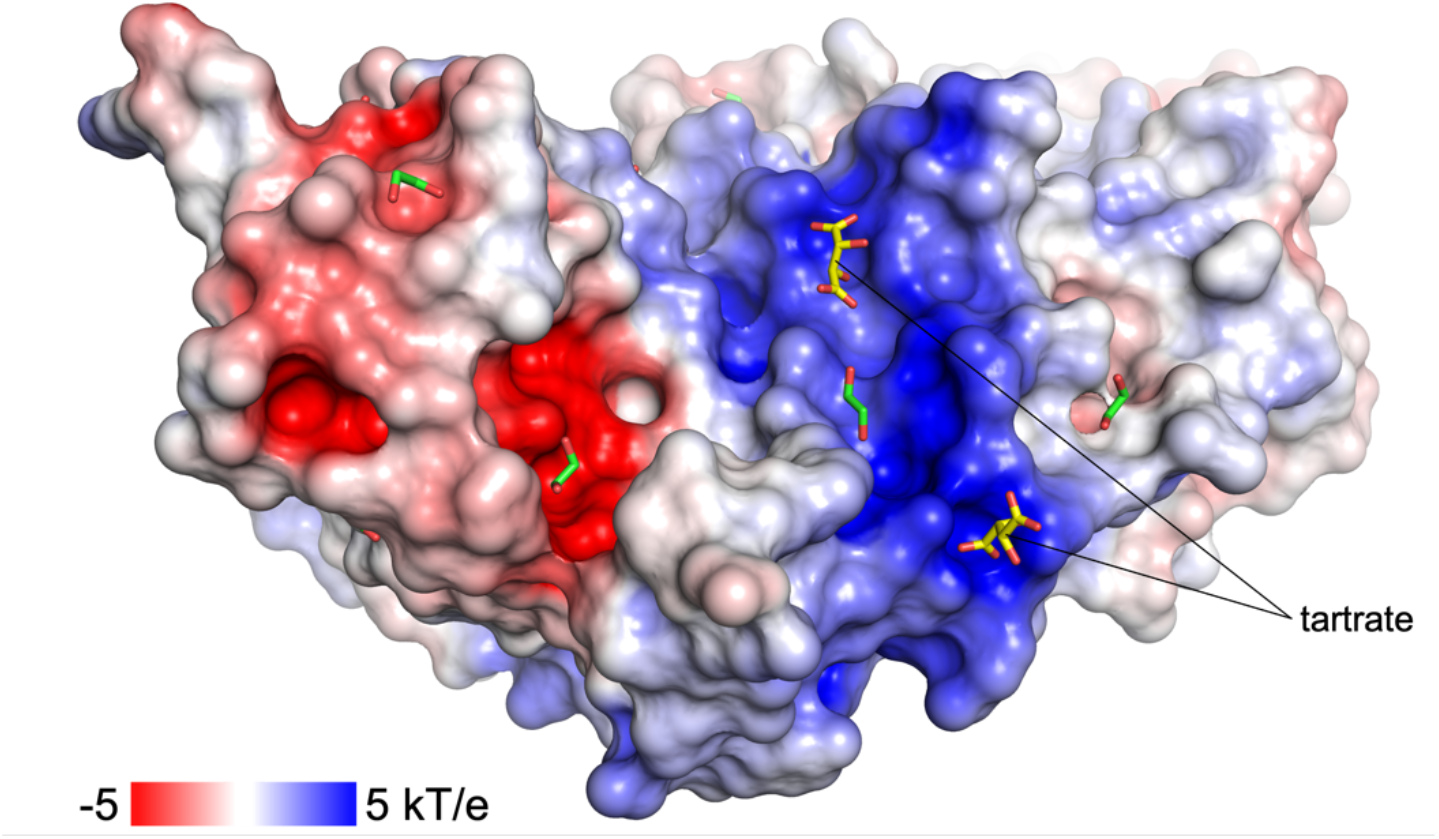
Tartrate ions bound on the basic patch of ExoN-nsp10 complex. Electrostatic surface potential of ExoN-nsp10 with tartrate or ethylene glycol bound on the protein surface.

**Supplementary Fig. 5.**
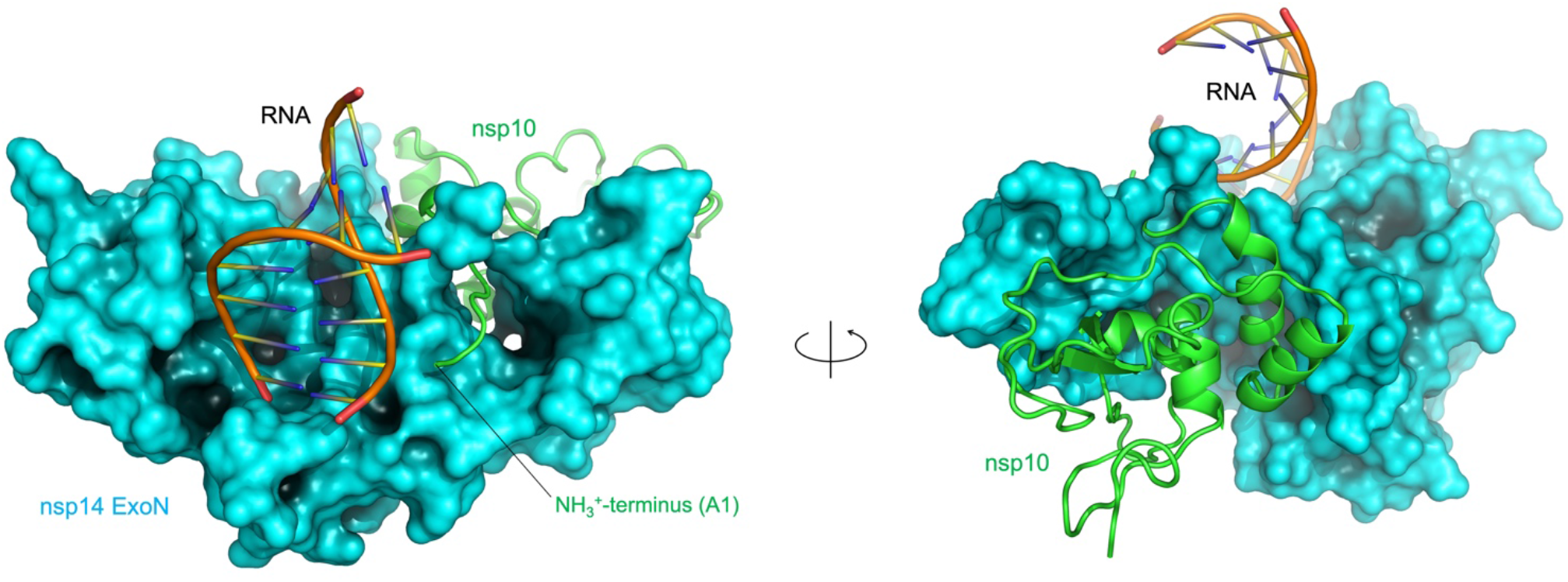
ExoN-nsp10-RNA complex model (an additional view) The image on the right is same as **Fig. 3C**.

**Supplementary Fig. 6.**
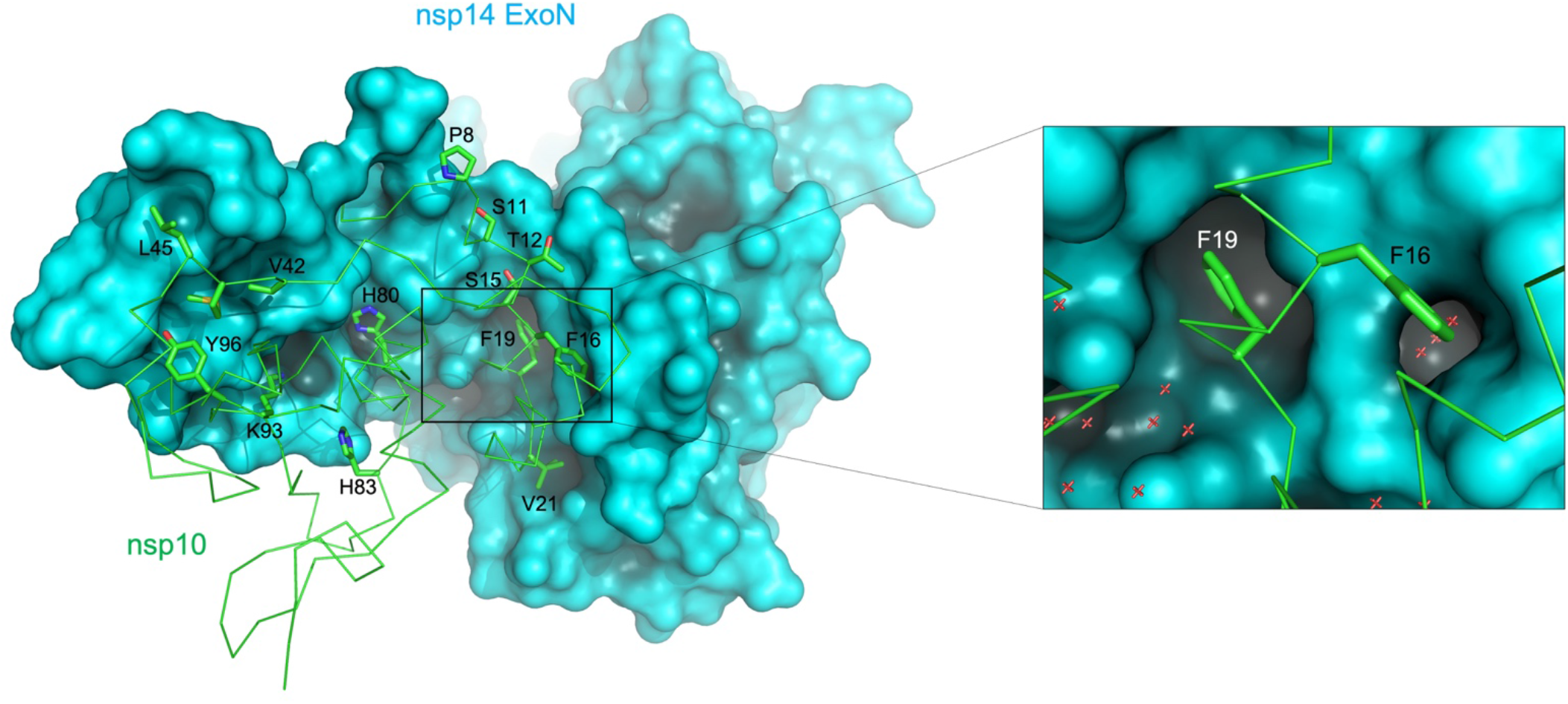
ExoN-nsp10 interface. ExoN is shown in solid surface and nsp10 in wire-frame representations, respectively. Some (not all) of the nsp10 side chains involved in the protein-protein interaction are shown as sticks. A zoomed view of the hydrophobic pocket that accommodates Phe16 and Phe19 of nsp10 is shown on the right. Red crosshairs represent water molecules.

**Supplementary Fig. 7.**
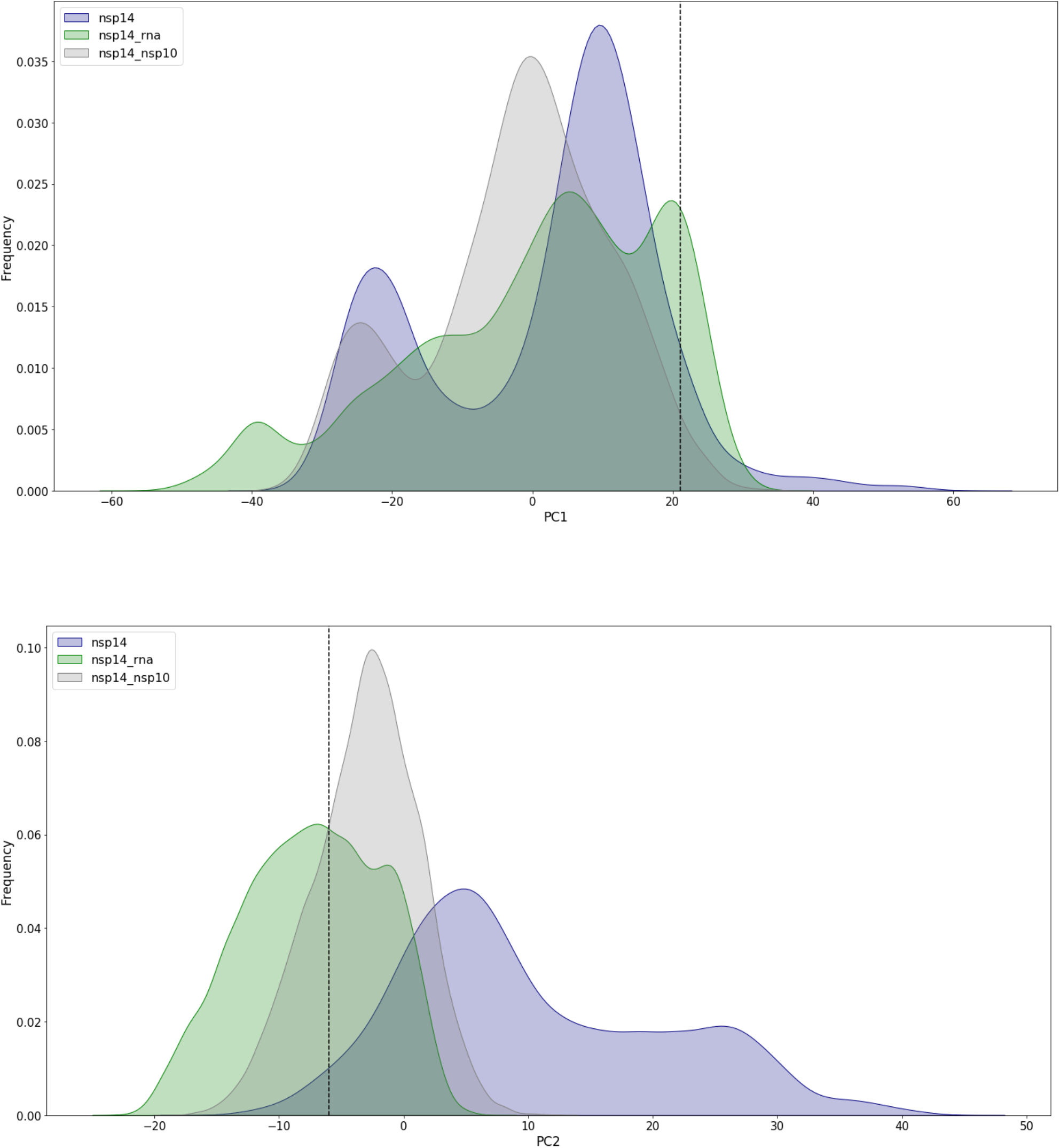
Distribution of principal components 1 and 2 (PC1 and PC2) observed in MD simulations. The dashed line in each plot indicates the value calculated for the starting structure.

**Supplementary Fig. 8.**
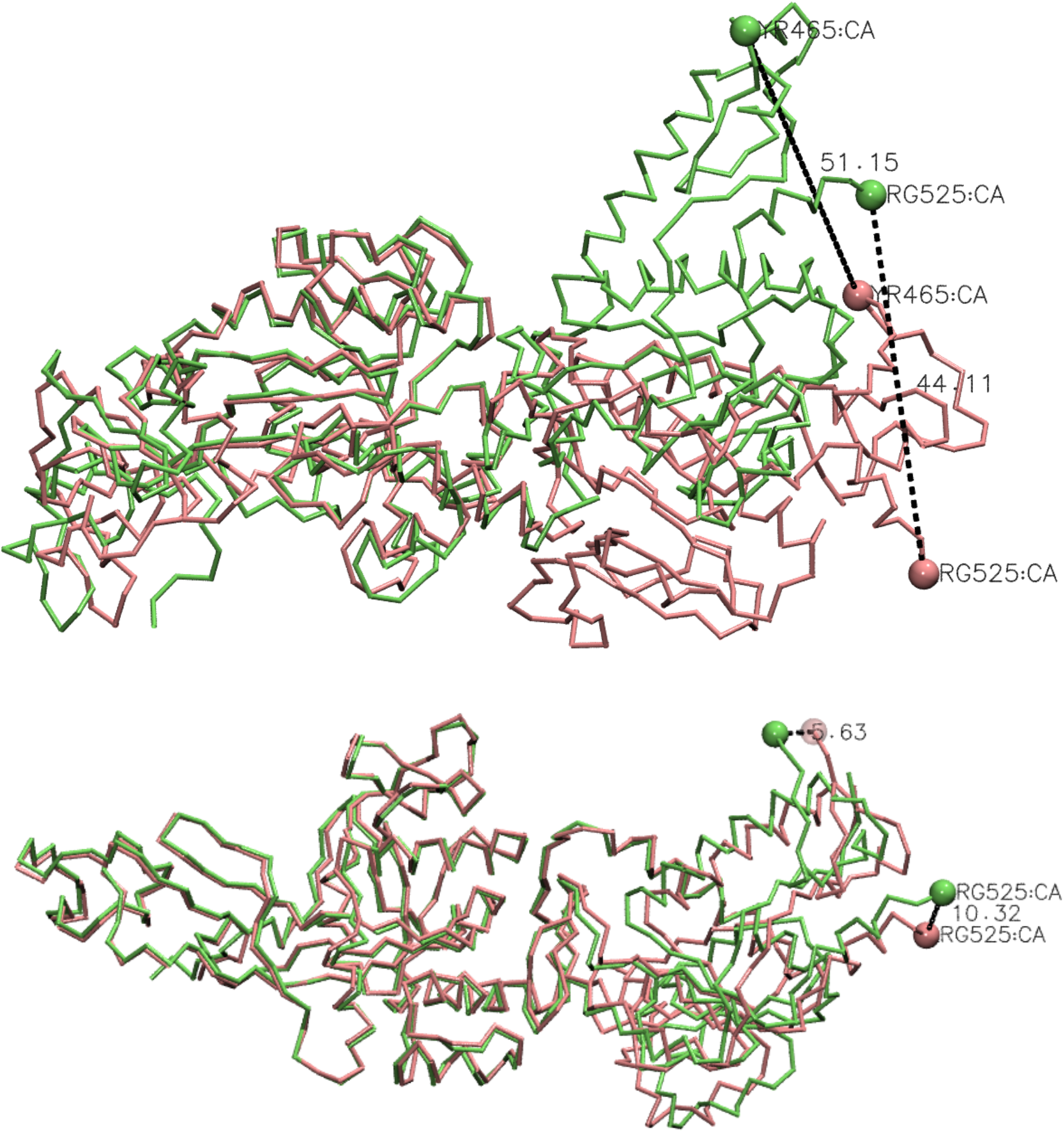
Conformational flexibility of full-length nsp14. **a**, Superposition of the structures that correspond to PC1 minimum and PC1 maximum values in MD simulations based on the ExoN domain, showing large displacement of the N7-MTase domain (This conformational change is also shown in **Supplementary animation 1)**. **b**, Similar conformational variability, albeit with a smaller magnitude, observed between chains A and B of SARS-CoV nsp14 crystal structure (PDB ID: 5NFY) (21).

**Supplementary Fig. 9.**
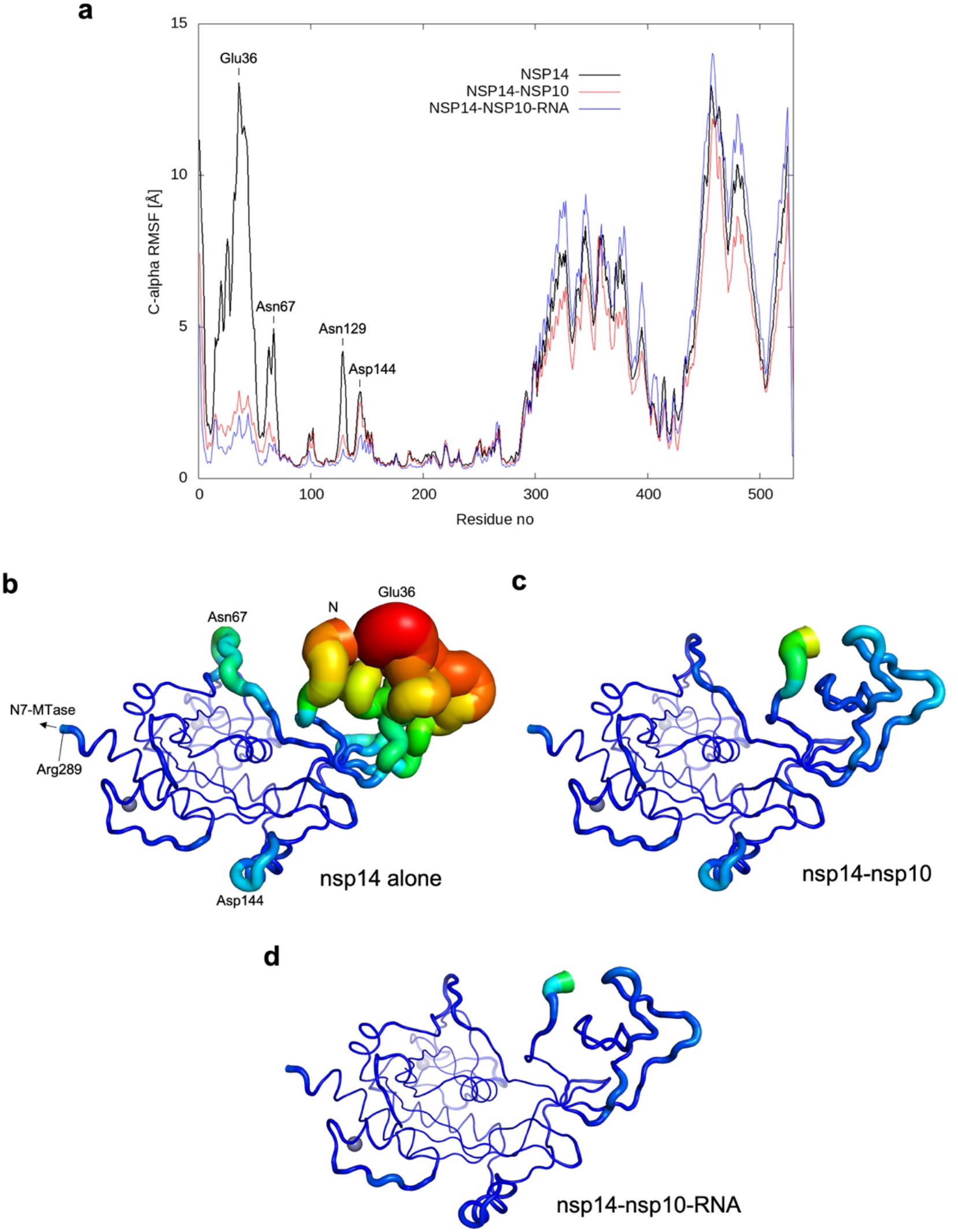
Internal dynamics of ExoN domain observed in MD simulations. **a**, Root-mean-square fluctuations (RMSF) for nsp14 Ca atoms in MD simulations of the three systems after aligning their trajectories to the starting structure with respect to Ca atoms of nsp14 residues 71-289. **b-d**, RMSF for nsp14 alone (b), nsp14-nsp10 (c), and nsp14-nsp10-RNA (d), depicted by tube thickness and color. Panels **b** and **c** are same as **Fig. 5 D** and **E**, respectively, and panels **b-d** correspond to the 3 frames in **Supplementary animation 3**.

**Supplementary Fig. 10.**
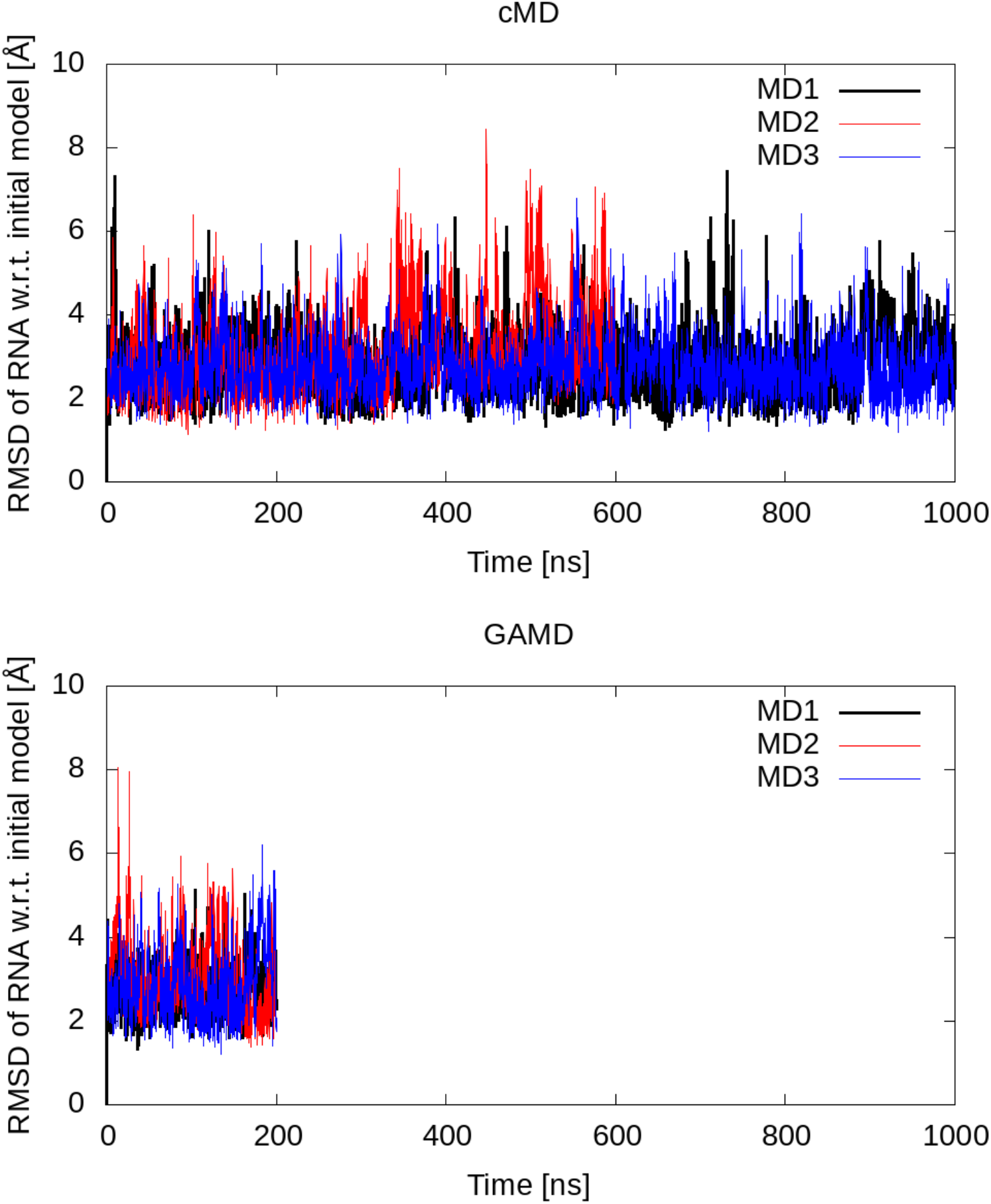
Stability of RNA in MD simulations. Root-mean-square deviation (RMSD) of RNA atoms calculated for conventional and Gaussian-accelerated MD (cMD and GAMD) simulations after aligning the trajectories with respect to Ca atoms of nsp14 residues 71-289.

**Supplementary Fig. 11.**
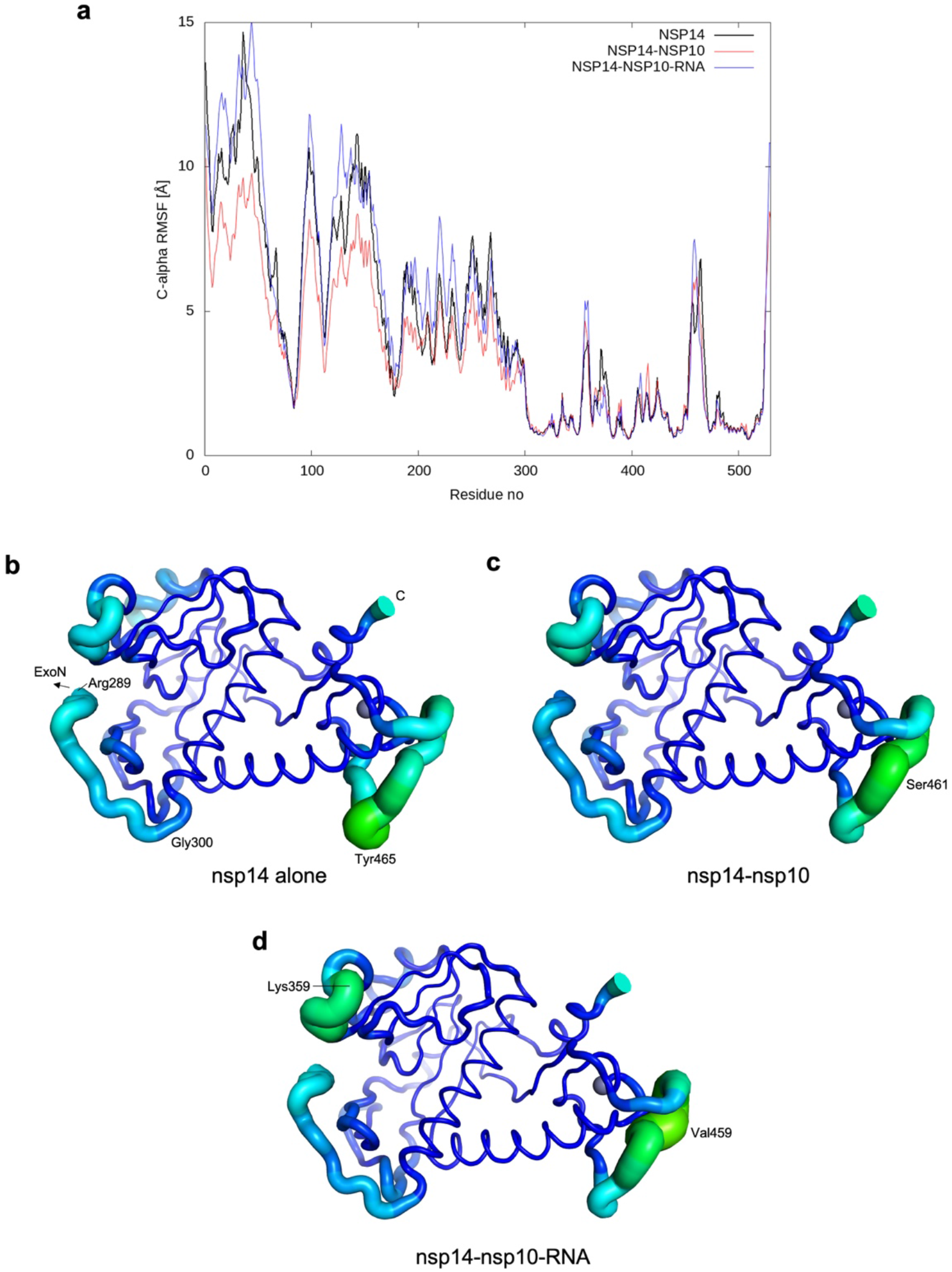
Internal dynamics of N7-MTase domain observed in MD simulations. **a**, Root-mean-square fluctuations (RMSF) for nsp14 Ca atoms in MD simulations of the three systems after aligning their trajectories to the starting structure with respect to Ca atoms of nsp14 residues 300-525. **b-d**, RMSF for nsp14 alone (b), nsp14-nsp10 (c), and nsp14-nsp10-RNA (d), depicted by tube thickness and color.

**Supplementary Fig. 12.**
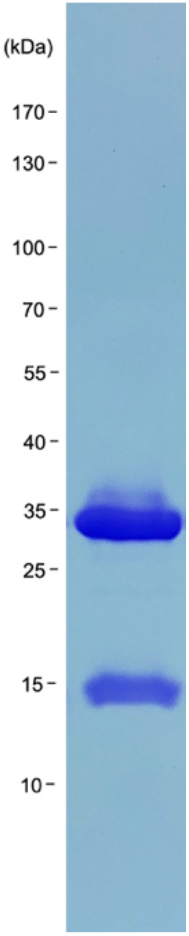
SDS-PAGE of purified SARS-CoV-2 ExoN(E191Q)-nsp10 complex. This protein complex was used in the crystallographic studies.

**Supplementary Fig. 13.**
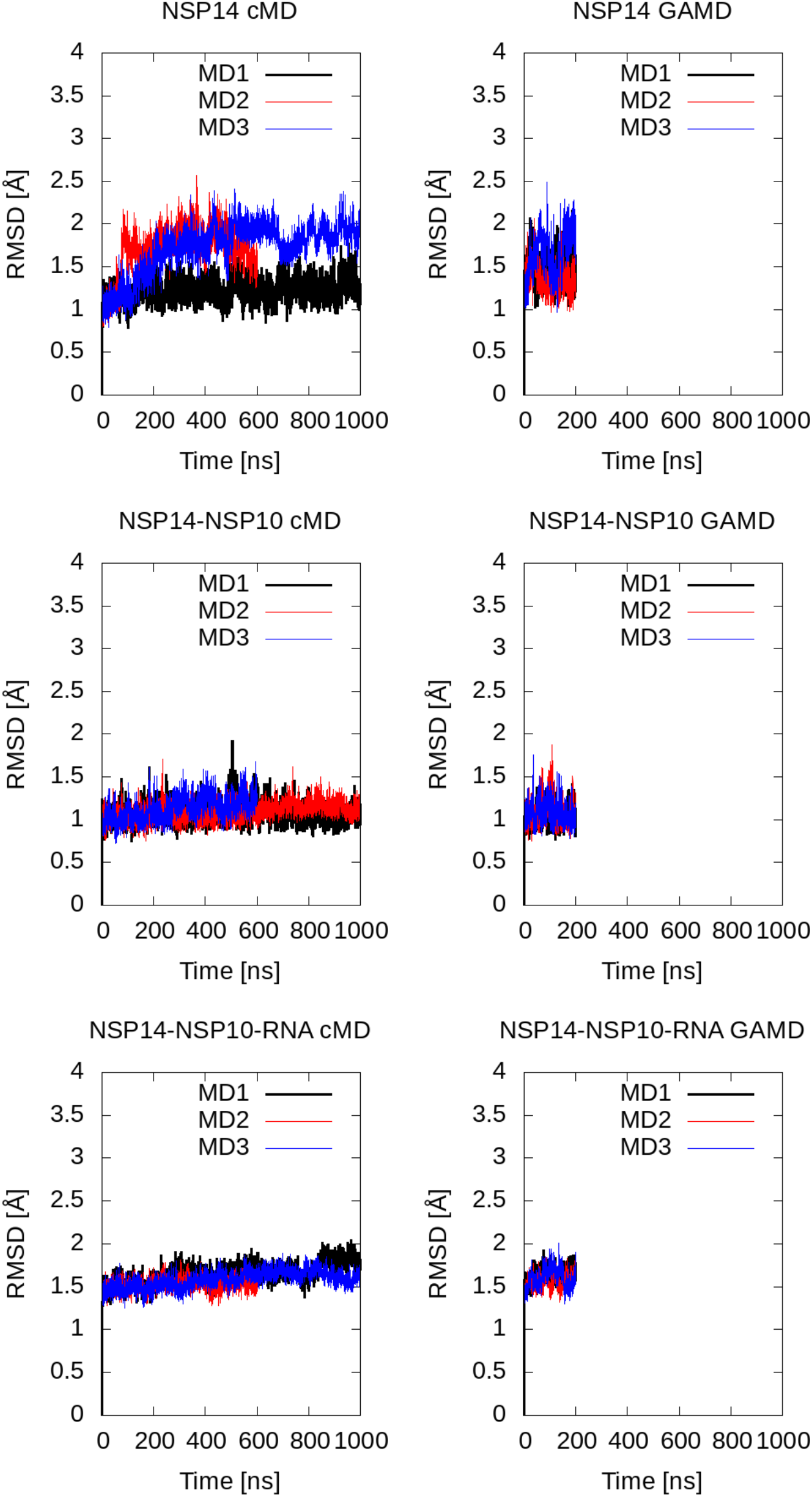
Stability of MD simulations. Root-mean-square deviation (RMSD) of the nsp14 ExoN domain Ca atoms (residues 71-289) with respect to the initial model throughout MD simulations.

## Reference

1. Hillen HS, et al. (2020 Structure of replicating SARS-CoV-2 polymerase. Nature 584(7819):154–156.

2. Posthuma CC, Te Velthuis AJW, & Snijder EJ (2017 Nidovirus RNA polymerases: Complex enzymes handling exceptional RNA genomes. Virus Res 234:58–73.

3. Sola I, Almazan F, Zuniga S, & Enjuanes L (2015 Continuous and Discontinuous RNA Synthesis in Coronaviruses. Annu Rev Virol 2(1):265–288.

4. Drake JW & Holland JJ (1999 Mutation rates among RNA viruses. Proc Natl Acad Sci U S A 96(24):13910–13913.

5. Jenkins GM, Rambaut A, Pybus OG, & Holmes EC (2002 Rates of molecular evolution in RNA viruses: a quantitative phylogenetic analysis. J Mol Evol 54(2):156–165.

6. Sanjuan R, Nebot MR, Chirico N, Mansky LM, & Belshaw R (2010 Viral mutation rates. J Virol 84(19):9733–9748.

7. Denison MR, Graham RL, Donaldson EF, Eckerle LD, & Baric RS (2011 Coronaviruses: an RNA proofreading machine regulates replication fidelity and diversity. RNA Biol 8(2):270–279.

8. Gorbalenya AE, Enjuanes L, Ziebuhr J, & Snijder EJ (2006 Nidovirales: evolving the largest RNA virus genome. Virus Res 117(1):17–37.

9. Shannon A, et al. (2020 Rapid incorporation of Favipiravir by the fast and permissive viral RNA polymerase complex results in SARS-CoV-2 lethal mutagenesis. Nat Commun 11(1):4682.

10. Minskaia E, et al. (2006 Discovery of an RNA virus 3′->5′ exoribonuclease that is critically involved in coronavirus RNA synthesis. Proc Natl Acad Sci U S A 103(13):5108–5113.

11. Robson F, et al. (2020 Coronavirus RNA Proofreading: Molecular Basis and Therapeutic Targeting. Mol Cell 79(5):710–727.

12. Smith EC & Denison MR (2013 Coronaviruses as DNA wannabes: a new model for the regulation of RNA virus replication fidelity. PLoS Pathog 9(12):e1003760.

13. Eskier D, Suner A, Oktay Y, & Karakulah G (2020 Mutations of SARS-CoV-2 nsp14 exhibit strong association with increased genome-wide mutation load. PeerJ 8:e10181.

14. Takada K, Takahashi Ueda M, Watanabe T, & Nakagawa S (2020 Genomic diversity of SARS-CoV-2 can be accelerated by a mutation in the nsp14 gene. bioRxiv 2020.12.23.424231; doi: https://doi.org/10.1101/2020.12.23.424231

15. Eckerle LD, et al. (2010 Infidelity of SARS-CoV Nsp14-exonuclease mutant virus replication is revealed by complete genome sequencing. PLoS Pathog 6(5):e1000896.

16. Eckerle LD, Lu X, Sperry SM, Choi L, & Denison MR (2007 High fidelity of murine hepatitis virus replication is decreased in nsp14 exoribonuclease mutants. J Virol 81(22):12135–12144.

17. Graham RL, et al. (2012 A live, impaired-fidelity coronavirus vaccine protects in an aged, immunocompromised mouse model of lethal disease. Nat Med 18(12):1820–1826.

18. Ogando NS, et al. (2020 The Enzymatic Activity of the nsp14 Exoribonuclease Is Critical for Replication of MERS-CoV and SARS-CoV-2. J Virol 94(23).

19. Gribble J, et al. (2021 The coronavirus proofreading exoribonuclease mediates extensive viral recombination. PLoS Pathog 17(1):e1009226.

20. Case JB, et al. (2018 Murine Hepatitis Virus nsp14 Exoribonuclease Activity Is Required for Resistance to Innate Immunity. J Virol 92(1).

21. Ferron F, et al. (2018 Structural and molecular basis of mismatch correction and ribavirin excision from coronavirus RNA. Proc Natl Acad Sci U S A 115(2):E162–E171.

22. Agostini ML, et al. (2018 Coronavirus Susceptibility to the Antiviral Remdesivir (GS-5734) Is Mediated by the Viral Polymerase and the Proofreading Exoribonuclease. mBio 9(2).

23. Smith EC, Blanc H, Surdel MC, Vignuzzi M, & Denison MR (2013 Coronaviruses lacking exoribonuclease activity are susceptible to lethal mutagenesis: evidence for proofreading and potential therapeutics. PLoS Pathog 9(8):e1003565.

24. Ma Y, et al. (2015 Structural basis and functional analysis of the SARS coronavirus nsp14-nsp10 complex. Proc Natl Acad Sci U S A 112(30):9436–9441.

25. Beese LS & Steitz TA (1991 Structural basis for the 3′-5′ exonuclease activity of Escherichia coli DNA polymerase I: a two metal ion mechanism. EMBO J 10(1):25–33.

26. Chen P, et al. (2007 Biochemical characterization of exoribonuclease encoded by SARS coronavirus. J Biochem Mol Biol 40(5):649–655.

27. Hastie KM, King LB, Zandonatti MA, & Saphire EO (2012 Structural basis for the dsRNA specificity of the Lassa virus NP exonuclease. PLoS One 7(8):e44211.

28. Jiang X, et al. (2013 Structures of arenaviral nucleoproteins with triphosphate dsRNA reveal a unique mechanism of immune suppression. J Biol Chem 288(23):16949–16959.

29. Bouvet M, et al. (2012 RNA 3′-end mismatch excision by the severe acute respiratory syndrome coronavirus nonstructural protein nsp10/nsp14 exoribonuclease complex. Proc Natl Acad Sci U S A 109(24):9372–9377.

30. Bouvet M, et al. (2014 Coronavirus Nsp10, a critical co-factor for activation of multiple replicative enzymes. J Biol Chem 289(37):25783–25796.

31. Saramago M, et al. (2021 New targets for drug design: Importance of nsp14/nsp10 complex formation for the 3′-5′ exoribonucleolytic activity on SARS-CoV-2. bioRxiv 2021.01.07.425745; doi: https://doi.org/10.1101/2021.01.07.425745

32. Baddock HT, et al. (2020 Characterisation of the SARS-CoV-2 ExoN (nsp14ExoN-nsp10) complex: implications for its role in viral genome stability and inhibitor identification. bioRxiv 2020.08.13.248211; doi: https://doi.org/10.1101/2020.08.13.248211.

33. Beese LS, Derbyshire V, & Steitz TA (1993 Structure of DNA polymerase I Klenow fragment bound to duplex DNA. Science 260(5106):352–355.

34. Chen Y, et al. (2013 Structure-function analysis of severe acute respiratory syndrome coronavirus RNA cap guanine-N7-methyltransferase. J Virol 87(11):6296–6305.

35. Kabsch W (2010 Xds. Acta Crystallogr D Biol Crystallogr 66(Pt 2):125–132.

36. McCoy AJ, et al. (2007 Phaser crystallographic software. J Appl Crystallogr 40(Pt 4):658–674.

37. Emsley P, Lohkamp B, Scott WG, & Cowtan K (2010 Features and development of Coot. Acta Crystallogr D Biol Crystallogr 66(Pt 4):486–501.

38. Liebschner D, et al. (2019 Macromolecular structure determination using X-rays, neutrons and electrons: recent developments in Phenix. Acta Crystallogr D Struct Biol 75(Pt 10):861–877.

39. Schrödinger (2021) Schrödinger Release 2021-1: Prime, Schrödinger, LLC, New York, NY.

40. Dolinsky TJ, Nielsen JE, McCammon JA, & Baker NA (2004 PDB2PQR: an automated pipeline for the setup of Poisson-Boltzmann electrostatics calculations. Nucleic Acids Res 32(Web Server issue):W665–667.

41. Maier JA, et al. (2015 ff14SB: Improving the Accuracy of Protein Side Chain and Backbone Parameters from ff99SB. J Chem Theory Comput 11(8):3696–3713.

42. Pang YP (1999 Novel Zinc Protein Molecular Dynamics Simulations: Steps Toward Antiangiogenesis for Cancer Treatment. J Mol Model 5:196–202.

43. Phillips JC, et al. (2020 Scalable molecular dynamics on CPU and GPU architectures with NAMD. J Chem Phys 153(4):044130.

44. Case DA, et al. (2020) AMBER 2020, University of California, San Francisco.

45. McGibbon RT, et al. (2015 MDTraj: A Modern Open Library for the Analysis of Molecular Dynamics Trajectories. Biophys J 109(8):1528–1532.

46. Baker NA, Sept D, Joseph S, Holst MJ, & McCammon JA (2001 Electrostatics of nanosystems: application to microtubules and the ribosome. Proc Natl Acad Sci U S A 98(18):10037–10041.

