## Supplementary animations for "Structure and dynamics of SARS-CoV-2 proofreading exoribonuclease ExoN"

**Supplementary animation 1**

Two nsp14 conformations corresponding to the minimum and maximum of principal component 1 **(**PC1) in MD simulations. The two frames are same as those in **Fig. 5B**.

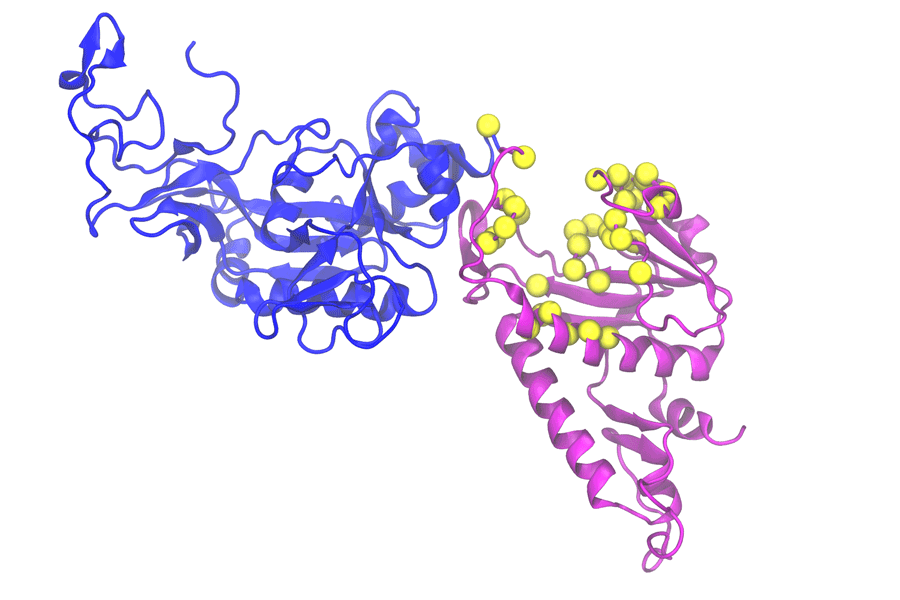

**Supplementary animation 2**

Two nsp14 conformations corresponding to the minimum and maximum of principal component 2 **(**PC2) in MD simulations. The two frames are same as those in **Fig. 5C**.

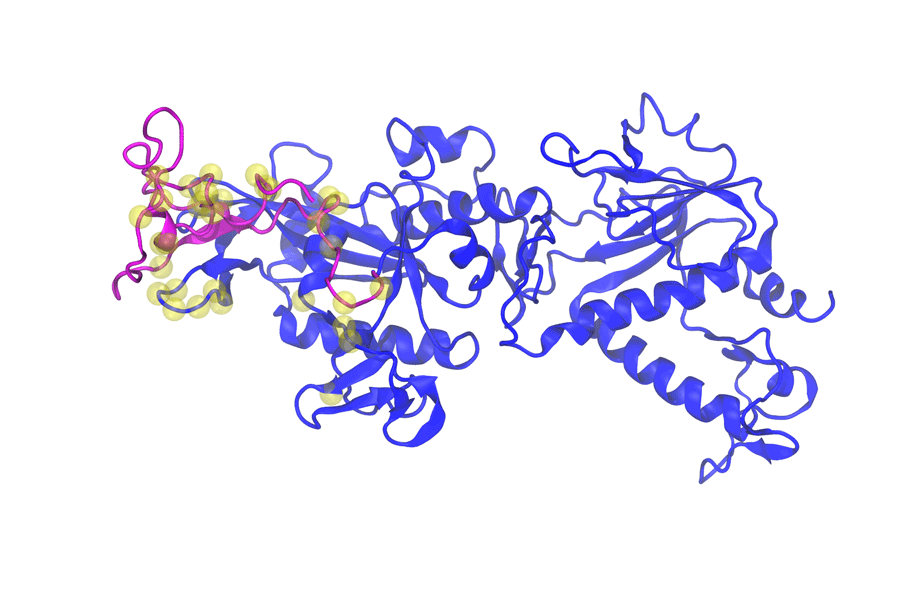

**Supplementary animation 3**

Root-mean-square fluctuations (RMSF) for nsp14 ExoN domain Cα atoms in MD simulations of the three systems (nsp14 alone, nsp14-nsp10, and nsp14-nsp10-RNA) after aligning their trajectories to the starting structure with respect to Cα atoms of nsp14 residues 71-289. RMSF is depicted by varying tube thickness and color. The 3 frames correspond to **Supplementary Fig. 9 b-d.**

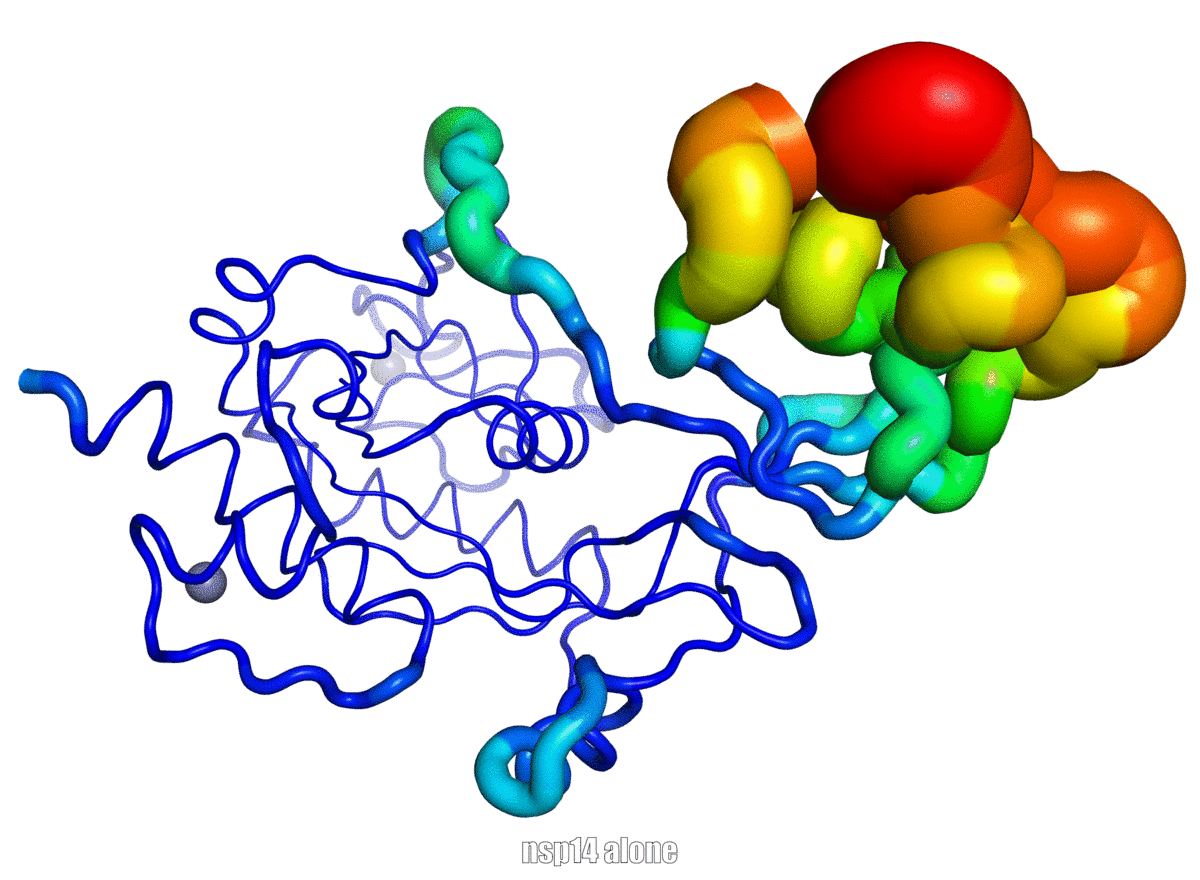
